# *LMX1B* missense-perturbation of regulatory element footprints disrupts serotonergic forebrain axon arborization

**DOI:** 10.1101/2024.12.12.628165

**Authors:** Brent Eastman, Nobuko Tabuchi, Xinrui Zhang, William C. Spencer, Evan S. Deneris

## Abstract

Pathogenic coding mutations are prevalent in human neuronal transcription factors (TFs) but how they disrupt development is poorly understood. Lmx1b is a master transcriptional regulator of postmitotic *Pet1* neurons that give rise to mature serotonin (5-HT) neurons; over two hundred pathogenic heterozygous mutations have been discovered in human *LMX1B,* yet their impact on brain development has not been investigated. Here, we developed mouse models with different *LMX1B* DNA-binding missense mutations. Missense heterozygosity broadly altered *Pet1* neuron transcriptomes, but expression changes converged on axon and synapse genes. Missense heterozygosity effected highly specific deficits in the postnatal maturation of forebrain serotonin axon arbors, primarily in the hippocampus and motor cortex, which was associated with spatial memory defects. Digital genomic footprinting (DGF) revealed that missense heterozygosity caused complete loss of Lmx1b motif protection and chromatin accessibility at sites enriched for a distal active enhancer/active promoter histone signature and homeodomain binding motifs; at other bound Lmx1b motifs, varying levels of losses, gains or no change in motif binding and accessibility were found. The spectrum of footprint changes was strongly associated with synapse and axon genes. Further, Lmx1b missense heterozygosity caused wide disruption of Lmx1b-dependent GRNs comprising diverse TFs expressed in *Pet1* neurons. These findings reveal an unanticipated continuum of Lmx1b missense-forced perturbations on *Pet1* neuron regulatory element TF binding and accessibility. Our work illustrates the power of DGF for gaining unique insight into how TF missense mutations interfere with developing neuronal GRNs.

**Significance Statement:** We modeled human LMX1B missense mutations in mice to explore how they disrupt brain serotonin neuron development. Missense heterozygosity selectively impaired postnatal formation of serotonin axon arbors throughout the forebrain, notably in the hippocampus and motor cortex. DGF revealed that Lmx1b missense heterozygosity exerted a continuum of footprint changes associated with synapse and axon gene expression. Footprint changes ranged from total eliminations to partial losses and gains within the *Pet1* neuronal epigenome. LMX1B missense mutations may cause human brain pathogenesis by selectively disrupting cis regulatory elements controlling 5-HT axon arbor formation thus impairing 5-HT delivery to presynaptic release sites.

## Introduction

Transcription factors (TFs) control the assembly of transcriptomes and epigenomes to establish specialized cell-type identities. Heterozygous coding mutations are prevalent in human transcription factor genes that are expressed in the human brain and can cause or increase risk for many neurodevelopmental disorders (1–4). Yet, few of these mutations have been investigated in an in vivo model system suitable for revealing how they may alter specific stages of neuronal development to cause brain pathogenesis. Further, a poorly studied question, particularly for mutated TF proteins that retain expression of functional domains, is how heterozygous TF mutations disrupt chromatin accessibility at regulatory elements required for neuronal gene expression.

The LIM homeodomain transcription factor, Lmx1b, is expressed in many developing and mature neuron-types in the brain (5, 6). Lmx1b together with Pet1 are key TFs in the regulatory program that generates brain serotonin (5-HT) neurons (7, 8). Lmx1b and Pet1 act in postmitotic *Pet1* neurons to induce expression of 5-HT neurotransmission genes required for brain 5-HT synthesis, reuptake, vesicular transport, and metabolism (9–12). These TFs also function at different developmental stages to control embryonic growth of long distance 5-HT projection pathways and postnatal 5-HT axon arborization in the brain and spinal cord, thereby enabling expansive serotonergic neuromodulation (13). For example, homozygous targeting of *Lmx1b* in newly born embryonic *Pet1* neurons results in failed development of 5-HT axon selective projections to the forebrain; homozygous targeting at different early postnatal stages, after secondary pathways have fully formed, result in severe deficits in 5-HT axon arborization (13). Their control of 5-HT neuron development involves reorganizing chromatin accessibility of cis-regulatory elements (CREs) associated with genes required for serotonergic neurotransmission, long distance axon growth and synapse formation (14). Further, Lmx1b is required for maintenance of adult 5-HT neurons as adult stage targeting of *Lmx1b* results in complete loss of expression of 5-HT neurotransmission genes and widespread progressive degeneration of 5-HT synapses and axons (15).

About 200 de novo or inherited heterozygous missense, nonsense, and deletion mutations have been described in the human LIM homeodomain transcription factor LMX1B (16–18). Most of the missense and nonsense mutations are clustered in the DNA-binding homeodomain (HD) and the dual LIM domains, which engage in protein-protein interactions. These mutations, many of which are recurrent, are pathogenic for a syndromic autosomal dominant disorder characterized by skeletal dysplasia, progressive renal disease, and glaucoma (16, 18–20). LMX1B haploinsufficiency is considered the underlying pathogenic mechanism (17). Intra- and interfamily heterogeneity in disease severity and penetrance is well established, yet there is no clear relation between mutation-type and clinical disease phenotype to account for clinical heterogeneity (16).

None of the pathogenic LMX1B mutations have been modeled to determine how they disrupt the development of the cell types in which they are expressed. Here, we developed new mouse lines carrying different recurrent pathogenic LMX1B DNA binding domain missense mutations. We leveraged the well-defined roles, summarized above, for Lmx1b in 5-HT neurons to assess how heterozygous Lmx1b missense mutations impact their development. We found that Lmx1b missense heterozygosity altered 5-HT neuron transcriptomes encoding synapse, axon, and mitochondrial processes. Further, missense heterozygosity resulted in a highly selective disruption of postnatal 5-HT axon arborization in the forebrain, which was associated with a sex-specific alteration in spatial memory formation. DGF revealed that Lmx1b missense heterozygosity exerted a complex combination of complete and partial losses as well as gains in Lmx1b motif binding and accessibility, associated with synapse and axon gene expression, in *Pet1* neuron chromatin. Lastly, Lmx1b missense heterozygosity caused wide disruption of Lmx1b-dependent gene regulatory networks (GRNs) comprising diverse TFs expressed in *Pet1* neurons. Our findings suggest LMX1B missense heterozygosity in humans may interfere with *Pet1* neuron cis regulatory element binding and accessibility required for control of postnatal forebrain 5-HT axon arbor formation thus compromising serotonergic neuromodulation.

## Results

### *LMX1B* missense mice

To generate missense line *Lmx1b^N246K^*, a C to A substitution identical to the human mutation was introduced in exon 5 changing an asparagine (N) codon to lysine (K) codon at amino acid position 246 of the mouse Lmx1b primary structure (Fig. 1A, Fig. S1A). This substitution is expected to result in a mutant protein unable to bind Lmx1b motifs, in vivo, as N246, an invariant residue in the HD recognition (third) helix makes direct hydrogen bond contacts with the second adenine in the HD binding core TAAT motif and the N246K substitution was reported to abolish Lmx1b motif binding in mobility shift assays (20–24). For mutant line, *Lmx1b^R200Q^*, a G to A substitution identical to the human mutation was introduced in exon 4 resulting in an arginine (R) to glutamine (Q) substitution at amino acid position 200 in the N-terminal arm of the HD (Fig. 1A, Fig. S1A). This mutation likely severely disables binding to Lmx1b motifs, in vivo, as the highly conserved R200 residue makes direct contact in the minor groove with the first thymine in the core TAAT motif and the R200Q mutation results in reduced Lmx1b motif binding, in vitro (19, 22–24). *Lmx1b^N246/+^* and *Lmx1b^R200Q/+^* mice are fully viable and capable of normal reproduction despite having reduced body weights (Fig. S1B). *Lmx1b^N246/+^* but not *Lmx1b^R200Q/+^* mice invariably exhibit an opaque eye pathology within the first six postnatal weeks (Fig. S1C).

**Fig. 1:**
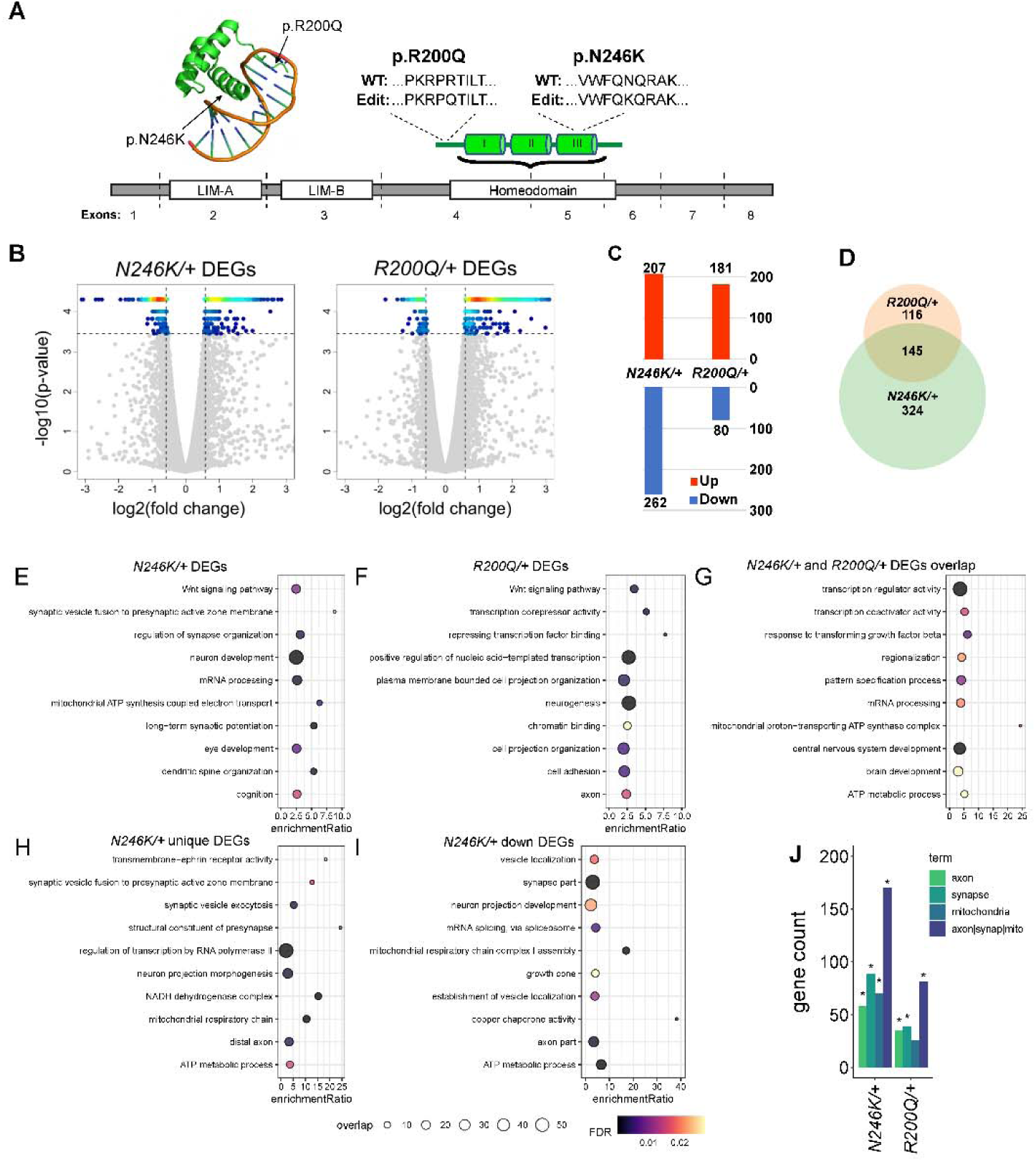
Altered development of *Lmx1b^N246K^* and *Lmx1b^R200Q^* heterozygous *Pet1* neuron transcriptomes. A) Top left, Model of HD third helix and N-terminal arm (green) binding to DNA. Arrows indicate approximate location of amino acid changes. Top right, sequence context of amino acid changes. Below, schematic of *Lmx1b* functional domains (white boxes) relative to exon boundaries. Green cylinders, HD alpha helices. B) Volcano plots displaying gene expression changes in *Lmx1b^N246K/+^* vs control and *Lmx1b^R200Q^*^/+^ vs control, with fold change ≥ 1.5X and FDR < 0.01 shown as -log10 p-value. Colored dots indicate significant DEGs. Dot heatmap indicates dot density: red, high; yellow/cyan, intermediate; blue, low. C) Number of up and down DEGs in *Lmx1b^N246K/+^* vs control and *Lmx1b^R200Q^*^/+^ vs control. D) Venn diagram of *Lmx1b^N246K/+^* and *Lmx1b^R200Q^*^/+^ common DEGs. E) *Lmx1b^N246K/+^* DEGs GO term enrichment. F) *Lmx1b^R200Q^*^/+^ DEGs GO term enrichment. G) *Lmx1b^N246K/+^* and *Lmx1b^R200Q^*^/+^ common DEGs GO term enrichment. H) *Lmx1b^N246K/+^* unique DEGs GO term enrichment. I) *Lmx1b^N246K/+^* down DEGs GO term enrichment J) Number of *Lmx1b^N246K/+^* and *Lmx1b^R200Q^*^/+^ DEGs that are annotated with axon, synapse, or mitochondria related GO terms. *, p = 0.00092 hypergeometric test, *Lmx1b^R200Q/+^* DEGs and p = 4.00E-12 hypergeometric test *Lmx1b^N246K/+^* DEGs. In E-I, dot size indicates the number of overlapping genes for each GO term and color indicates FDR.

No *Lmx1b^N246K^*^/*N246K*^ or *Lmx1b^R200Q/R200Q^* newborns survived beyond P1, consistent with previous neonatal lethality observed for homozygous *Lmx1b* targeted mice in which exons 3-7 encoding the second LIM domain and the HD were deleted (25). The tectum, cerebellum and hindbrain were severely reduced in size in *Lmx1b^N246K^*^/*N246K*^ and *Lmx1b^R200Q/R200Q^* newborns (Fig. S2A). Similar defects were found in germ line-conditionally targeted homozygous mice (*Lmx1b*^Δ*/*Δ^) in which the entire HD was removed to create a total loss of function DNA binding null. These findings further support the conclusion that missense protein binding to Lmx1b motifs is severely compromised or even eliminated, in vivo.

We introduced the *Pet1-Cre* transgene and *Ai9* allele into *Lmx1b^N246K^* and *Lmx1b^R200Q^* mice to genetically mark postmitotic *Pet1* neuron precursors from which brain 5-HT neurons develop (26). We found significant deficits in TdTomato^+^ cell body numbers at E15.5 in *Lmx1b^N246K/N246K; Pet1-Cre; Ai9^* and *Lmx1b*^Δ*/*Δ;^ *^Pet1-Cre; Ai9^* embryos (Fig. S2B, D), suggesting a previously unrecognized role for Lmx1b in the generation of *Pet1* neuron precursors prior to them becoming postmitotic. There was a trend for loss of TdTomato^+^ cell bodies in *Lmx1b^R200Q/R200Q; Pet1-Cre; Ai9^* mice but this did not reach statistical significance (Fig. S2B, D). A complete anterior to posterior analysis of the dorsal raphe nucleus (DRN), median raphe nucleus (MRN), B9 cluster, and medullary raphe nuclei revealed no significant differences in TdTomato^+^ cell body numbers in *Lmx1b^N246K/+; Pet1-Cre; Ai9^*, *Lmx1b^R200Q/+; Pet1-Cre; Ai9^* and *Lmx1b*^Δ^*^/+^*; *^Pet1-Cre; Ai9^* mice compared to controls (*Lmx1b^+/+; Pet1-Cre; Ai9^*) (Fig. S2C, E, F).

### Missense heterozygosity disrupts *Pet1* neuron synapse, axon, and mitochondrial transcriptomes

We performed RNA-seq to determine how *Lmx1b^N246K^* and *Lmx1b^R200Q^* missense heterozygosity impacts development of *Pet1* neuron transcriptomes. Sequencing libraries were prepared with flow sorted TdTomato^+^ *Lmx1b^N246K/+; Pet1-Cre; Ai9^* and *Lmx1b^R200Q/+; Pet1-Cre; Ai9^ Pet1* neurons taken from the entire E17.5 embryonic hindbrain to capture developing 5-HT neurons from all raphe nuclei. Differential expression analysis indicated that *Lmx1b^N246K^* and *Lmx1b^R200Q^* heterozygosity resulted in significant alterations in the development of *Pet1* neuron transcriptomes (Fig. 1B). We found 469 and 261 significantly increased or decreased differentially expressed genes (DEGs) with ≥1.5 fold change and ≤1% FDR in *Lmx1b^N246K/+; Pet1-Cre; Ai9^* and *Lmx1b^R200Q/+; Pet1-Cre; Ai9^ Pet1* neurons, respectively (Fig. 1C). 20% of DEGs (145/729) were found in both *Lmx1b^N246K/+^* and *Lmx1b^R200Q/+^* transcriptomes suggesting each missense protein forces mostly unique changes in *Pet1* neuron gene expression (Fig. 1D).

We used gene ontology enrichment analysis to gain insight into potential functions of genes with altered expression in *Lmx1b^N246K/+^*and *Lmx1b^R200Q/+^ Pet1* neurons. GO analysis of *Lmx1b^N246K/+^* or *Lmx1b^R200Q/+^* DEGs identified enrichment for several terms related to synapse, axon, and mitochondrial (SAM) genes such as synaptic vesicle fusion to presynaptic zone membrane, regulation of synapse organization, cell projection organization, axon, and mitochondrial ATP synthase coupled electron transport (Fig. 1E, F). The 145 common DEGs were enriched for terms related to transcription, mRNA processing, mitochondria function and brain development (Fig. 1G). Strong enrichment for SAM terms were also found when we analyzed *Lmx1b^N246K/+^*unique DEGs and *Lmx1b^N246K/+^* down DEGs (Fig. 1H, I). The log2 fold change (log2FC) range for SAM DEGs was −3X to 2X for *Lmx1b^N246K/+^* and −1.3X to 2.7X for *Lmx1b^R200Q/+^*. We found that 31% (81/261, p = 0.00092 hypergeometric test) of *Lmx1b^R200Q/+^* DEGs and 37% (171/469, p = 4.00E-12 hypergeometric test) of *Lmx1b^N246K/+^* DEGs were annotated with at least one of these classification terms (Fig. 1J). These findings indicate that both heterozygous missense mutations alter diverse types of genes that converge on processes related to development of serotonergic connectivity. Other enriched terms of lower significance did not converge on particular biological, cellular, or molecular categories despite finding that gene expression is more broadly altered than just changes in SAM expression (Fig. 1E-J). No term enrichment was found for processes related to development of 5-HT neuron identity.

5-HT levels and expression of all genes required for 5-HT neurotransmission are severely reduced in homozygous germ line or postmitotic stage conditionally targeted *Lmx1b* mice, (9, 11, 12, 27). Similarly, we found a complete absence of 5-HT immunoreactivity in the rostral ventral hindbrain in *Lmx1b^N246K^*^/*N246K*^ or *Lmx1b^R200Q/R200Q^* in E15.5 embryos (Fig. S3A) showing that both missense mutations produce equivalent loss of function alleles for acquisition of 5-HT neuron identity. Although expression of 5-HT synthesis genes, *Tph2, Ddc, Gch1, Gchfr*, trended down in *Lmx1b^R200Q/+^* mice the loss of expression was not significantly different from control levels (Fig. S3B). Only *Slc6a4* expression was significantly reduced by about 40% (−0.71X log2FC) in *Lmx1b^R200Q/+^ Pet1* neurons. *Lmx1b^N246K^*heterozygosity resulted in −0.75X and 0.6X log2FC decreased expression for *Tph2* and *Ddc,* respectively, but did not significantly alter expression of *Gch1* or *Gchfr*, which are required for 5-HT synthesis through production of the obligatory Tph2 co-factor, BH4 (Fig. S3B). *Pet1* expression is dramatically reduced in homozygous *Lmx1b* targeted mice, but its expression was not altered in *Lmx1b^N246K/+^*or *Lmx1b^R200Q/+^* mice (Fig. S3B). Although *Tph2* and *Ddc* expression was reduced, we found no alterations in the numbers of Tph2^+^ and 5-HT^+^ neurons in adult *Lmx1b^N246K/+^*and *Lmx1b^R200Q/+^*mice and individual neuron staining intensities seemed invariant between the three genotypes (Fig. S3C-F). These findings suggest that one wildtype copy of *Lmx1b* is sufficient for generation of normal numbers of 5-HT neuron postmitotic precursors and decreased *Tph2* and *Ddc* expression was not sufficient to weaken immunohistochemical detection of 5-HT neuron identity markers.

Homozygous conditional targeting of *Lmx1b* revealed its role in formation of the isthmic organizer and in the specification of progenitors that likely give rise to DA neurons in the substantia nigra pars compacta (SNc) (28–30). However, immunostaining with an anti-tyrosine hydroxylase (TH) antibody revealed no differences in the numbers of TH^+^ neurons in the SNc and TH^+^ fiber densities in the dorsal striatum of heterozygous mutant mice compared to controls (Fig. S4).

### Forebrain 5-HT axon deficits in *Lmx1b^N246K^* and *Lmx1b^R200Q^* heterozygotes

Given the strong convergence of *Lmx1b^N246K/+^* or *Lmx1b^R200Q/+^* DEGs on SAM genes, we investigated whether *Lmx1b^N246K^* and *Lmx1b^R200Q^* heterozygosity disrupted 5-HT axon development. Our previous studies showed that homozygous conditional targeting of the *Lmx1b* (*Lmx1b^f/f;Pet1-Cre;Ai9^*) specifically in postmitotic embryonic *Pet1* neurons resulted in near complete absence of 5-HT axons in the distal forebrain and spinal cord (13). Notably, although not as severe as that found in *Lmx1b^f/f;Pet1-Cre;Ai9^*mice, we found significantly reduced TdTomato^+^ fiber densities in virtually all regions of the forebrain in *Lmx1b^N246K/+; Pet1-Cre; Ai9^* and *Lmx1b^R200Q/+; Pet1-Cre; Ai9^* mice. To facilitate accurate quantification of axon deficits, we focused on regions receiving dense serotonergic innervation. In the hippocampus and motor cortex, comparable reductions in TdTomato^+^ fiber densities were observed in *Lmx1b^N246K/+; Pet1-Cre; Ai9^*and *Lmx1b^R200Q/+; Pet1-Cre; Ai9^* mice compared to control (Fig. 2A-E). However, TdTomato^+^ fibers in the striatum and the dense plexus in subventricular zone (SVZ), were significantly reduced in *Lmx1b^N246K/+; Pet1-Cre; Ai9^* mice but not in *Lmx1b^R200Q/+; Pet1-Cre; Ai9^* mice (Fig. S5A-D), a difference which may result from the large number of non-overlapping DEGs in the mutant transcriptomes. TdTomato^+^ fiber densities in the spinal cord were not statistically different from control in missense heterozygotes although we found a statistically significant difference in *Lmx1b*^Δ*/+;*^ *^Pet1-Cre; Ai9^* versus control (Fig. S6A, B).

**Figure 2:**
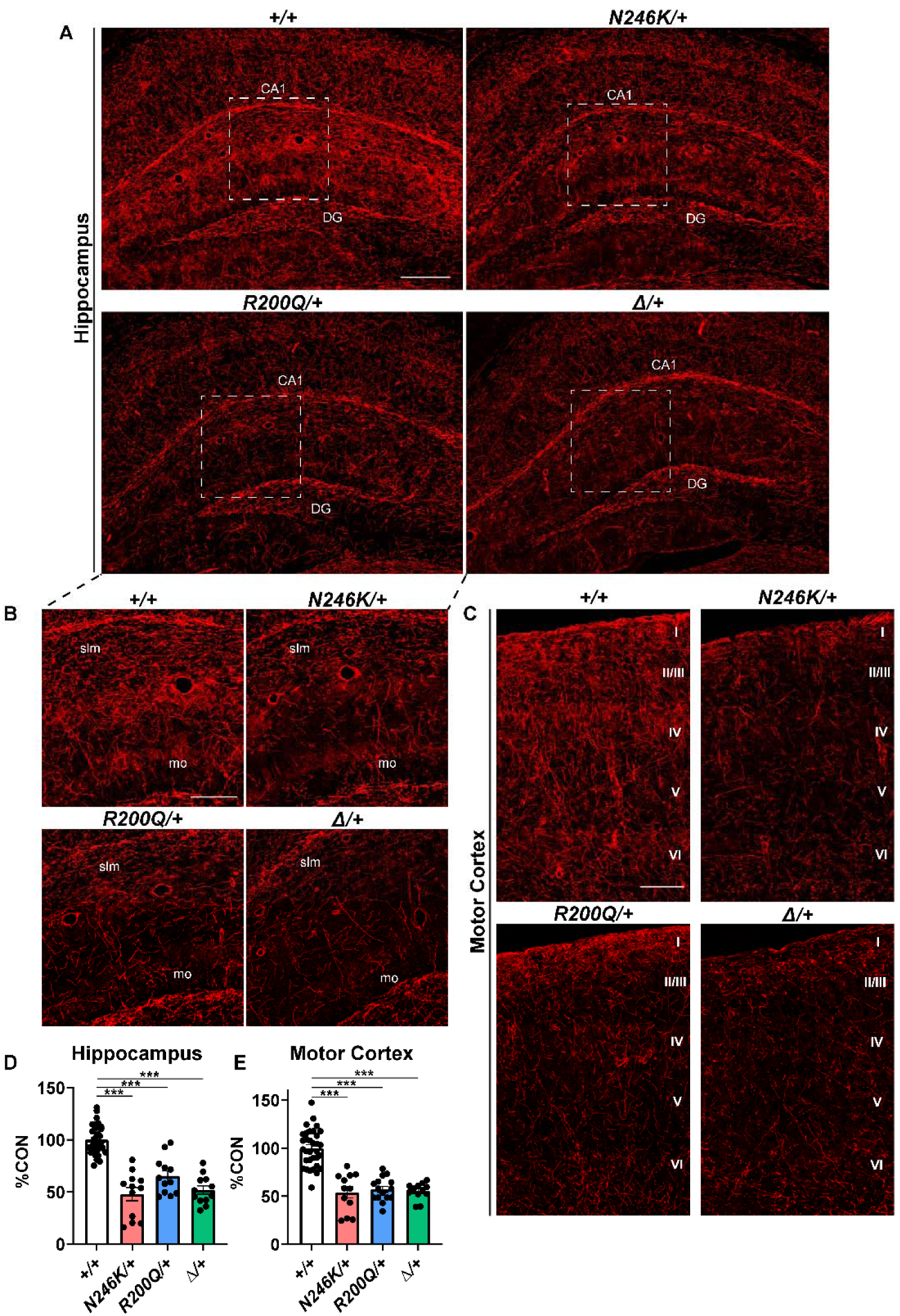
Forebrain serotonergic axon deficits in heterozygous adult missense mice. A) Representative images of TdTomato^+^ (anti-RFP) axons in the adult hippocampus. Scale bar, 200µm. DG, dentate gyrus; CA1, hippocampal subfield cornu ammonis 1. B) Higher magnification images of highlighted regions (white dashed outline) in A. Scale bar, 100µm. slm, stratum lacunosum moleculare; mo, molecular layer. C) Representative images of TdTomato^+^ (anti-RFP) axons in the adult motor cortex. Scale bar, 100µm. D-E) Relative TdTomato^+^ (anti-RFP) pixel densities from three representative sections per biological replicate; ±SEM; n=4-10 mice per genotype; one-way ANOVA with Welch’s correction. ***p<0.001.

### *Lmx1b^N246K^* and *Lmx1b^R200Q^* heterozygosity selectively disrupts postnatal development of 5-HT axon arbors

Lmx1b is required to maintain adult 5-HT axons (15), and so we sought to determine whether the 5-HT axon deficits in *Lmx1b^N246K/+^* or *Lmx1b^R200Q/+^* mice resulted from their abnormal development or post-developmental degeneration. Development of rodent 5-HT axon architectures has been described as comprising three sequential stages: primary axon growth, secondary routing and arborization (13, 31). During the primary stage, which initiates in newborn postmitotic *Pet1* neurons before they have acquired a fully mature serotonergic identity, axons ascend unidirectionally over the mesencephalic flexure. The secondary routing stage occurs between E16-P1 and results in formation along preexisting non-serotonergic fiber tracts of all 5-HT fiber pathways to their termination zones in the forebrain. The arborization stage occurs in the postnatal period subsequent to secondary routing and is not complete until at least the fourth postnatal week (32).

Although a delay in primary axon growth was present in homozygous conditionally targeted *Lmx1b* (*Lmx1b^f/f;Pet1-Cre;Ai9^*) embryos (13), no delay was found in *Lmx1b^N246K/+^*or *Lmx1b^R200Q/+^* embryos (Fig. S7A, E). Development of serotonergic fibers through the cingulum bundle culminates in the formation of a major 5-HT system secondary route to the forebrain (33). In *Lmx1b^f/f;Pet1-Cre;Ai9^* mice, the cingulum fiber track fails to form (13). At E18.5, TdTomato^+^ fiber densities within the cingulum bundle were not altered in *Lmx1b^N246K/+; Pet1-Cre; Ai9^* and *Lmx1b^R200Q/+; Pet1-Cre; Ai9^* embryos indicating that full Lmx1b dosage is not required for development of this major serotonergic projection pathway (Fig. S7B, F). A similar lack of primary and secondary fiber deficits was found in *Lmx1b*^Δ*/+;*^ *^Pet1-Cre; Ai9^*embryos confirming that complete loss of sequence-specific DNA binding in one Lmx1b copy had no detectable impact on this stage of 5-HT axon development.

At P7, we found a significant reduction in TdTomato^+^ fiber densities in the motor cortex of *Lmx1b^N246K/+; Pet1-Cre; Ai9^* and *Lmx1b^R200Q/+; Pet1-Cre; Ai9^* mice (Fig. S7C, G). In one month old *Lmx1b^N246K/+; Pet1-Cre; Ai9^* and *Lmx1b^R200Q/+; Pet1-Cre; Ai9^* mice, we found significant reductions in TdTomato^+^ fiber densities in the hippocampus (Fig. S7D, H). Together, these data demonstrate that heterozygous Lmx1b missense mutations specifically disrupt the postnatal stage of 5-HT axon development during which profuse arborization occurs.

Previous studies have implicated dense serotonergic inputs to hippocampal CA1 circuitry at the interface of the stratum radiatum (SR) and the stratum lacunosum-moleculare (SLM) layers in the modulation of spatial memory formation (34). Given the deficit in 5-HT arbors in this area of the hippocampus of missense heterozygotes, we examined spatial memory in *Lmx1b^R200Q/+^* mice. We chose not to test *Lmx1b^N246K/+^* mice because of their eye pathology. In the Barnes Maze, both male and female *Lmx1b^R200Q/+^* mice found the escape hatch as quickly as *Lmx1b^+/+^* on the final day of training. However, during the probe trial female *Lmx1b^R200Q/+^* mice spent significantly less time in the target quadrant compared to sex matched controls (Fig. S8). No difference in the time spent in the target quadrant was found between male *Lmx1b^R200Q/+^* mice and controls although males exhibited reduced distance traveled in the open field (Fig. S8).

### Lmx1b heterozygous missense mutations force a complex combination of losses and gains in Lmx1b motif binding and accessibility

We next sought to understand how Lmx1b missense heterozygosity impacts Lmx1b binding activity within the *Pet1* neuron epigenome. We previously used CUT&RUN to map Lmx1b occupancy within wild type *Pet1* neuron chromatin (14). However, we anticipated that prohibitively large numbers of flow sorted control and mutant *Pet1* neurons would be required to discern differences in Lmx1b binding with CUT&RUN. Here, we selected DGF to investigate Lmx1b binding activity (35–38). DGF identifies regions of lower-than-expected transposase-mediated cut counts centered on specific DNA binding motifs, termed footprints, that result from steric protection by a motif bound TF. A footprint score can be calculated by subtracting the mean of negative signal at the footprint from the mean of positive signal representing chromatin accessibility in the immediate flanking region. Unlike CUT&RUN or ChIP-seq, DGF offers genome-wide information on TF binding activity at motifs comprising disparate GRNs and accessibility of CREs in which TF motifs are embedded.

Bentsen et al., applied a novel framework (TOBIAS) for correction of Tn5 cut bias to determine the developmental dynamics of TF footprints in ATAC-seq datasets (38). Therefore, we crossed the *Sun1-GFP* allele and the *Pet1-Cre* BAC transgene into wild type, *Lmx1b^N246K/+^*and *Lmx1b^R200Q/+^* mice for purification of GFP-labeled nuclei from *Pet1* neurons in the embryonic hindbrain with Isolation of Nuclei TAgged in specific Cell Types (INTACT) (39). Purified nuclei from the three genotypes were tagmented with an ATAC-seq protocol optimized for *Pet1* neurons; libraries were deeply sequenced, replicates merged, and reads downsampled to 100 million/genotype (40). Comparing read counts within CREs of *Lmx1b^+/+^*, *Lmx1b^N246K/+^,* and *Lmx1b^R200Q/+^* ATAC-seq datasets with previously obtained control *Pet1* neuron ATAC-seq datasets at the same time point showed high concordance (Pearson’s correlation = 0.86-0.87) indicating high dataset reproducibility (14). We used TF motifs from the JASPAR 2022 database, which includes the human LMX1B motif derived with the HT-SELEX method (41).

Of the 23,744 total Lmx1b motifs identified, TOBIAS identified 4,493 bound footprints indicating likely sites of Lmx1b occupancy in control *Pet1* neuron chromatin. These footprints were widely distributed across the *Pet1* epigenome but were more abundant in introns and distal intergenic regions than promoters (Fig. S9A). The aggregate footprint profile showed strong protection at Lmx1b motifs and high accessibility in the immediate flanking regions (Fig. 3A, Fig. S9B). A significant fraction of Lmx1b-bound footprints overlaps with Lmx1b CUT&RUN peaks in *Pet1* neurons (permutation test p-value < 0.001, Fig. S9C). Additionally, a significant fraction of Pet1-bound footprints overlaps with Pet1 CUT&RUN peaks (p-value < 0.001) and CTCF-bound footprints overlap with CTCF ChIPmentation peaks (p-value < 0.001) (Fig. S9C). These data indicate the TOBIAS footprinting data reflects TF binding in vivo. When intersecting bound footprints, we found that less than half of control bound footprints were called bound in the missense heterozygous *Pet1* neurons (Fig. S9D) resulting from losses of bound footprints in the mutants. The aggregate footprint profile across all bound Lmx1b motifs was decreased in *Lmx1b^N246K/+^*and *Lmx1b^R200Q/+^ Pet1* neurons compared to control indicating overall reduced Lmx1b motif binding activity and loss of flanking accessibility (Fig. 3A, Fig. S9B). However, the *Lmx1b^N246K^* mutation resulted in a more severely decreased aggregate footprint score (−0.47X log2FC, p-value = 4.30E-181) than *Lmx1b^R200Q^* (−0.29X log2FC, p-value = 3.57E-160), indicating that similar to their impact on 5-HT neuron axon arbors and transcriptomes, the impact of the two HD missense mutations on the *Pet1* neuron epigenome are not equivalent.

**Figure 3:**
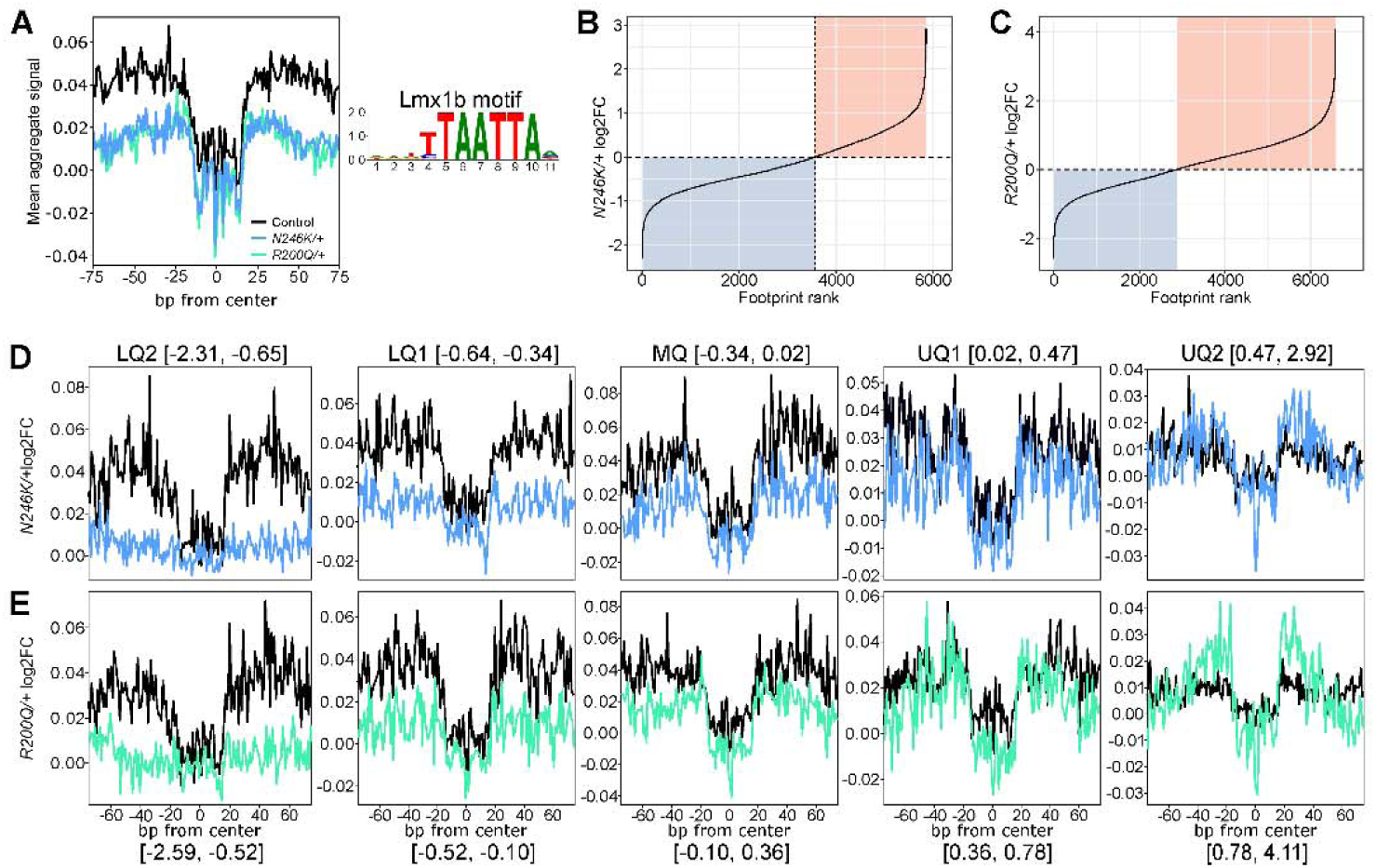
L*m*x1b heterozygous missense mutations disrupt Lmx1b binding activity and flanking chromatin accessibility. A) Aggregate footprint profiles for Lmx1b motif binding activity in control (black), *Lmx1b^N246K/+^* (green), and *Lmx1b^R200Q^*^/+^ (blue) *Pet1* neurons. Right, Lmx1b position weight matrix binding motif. B-C) Log2FC plots of Lmx1b motif footprint scores for *Lmx1b^N246K/+^* (B) and *Lmx1b^R200Q^*^/+^ (C) *Pet1* neurons. Colored quadrants highlight proportion of footprint losses (blue) and footprint gains (red) with greater proportion of losses in *N246K/+* vs *R200Q/+* and greater gains in *R200Q/+*. D-E) *Lmx1b^N246K/+^* (top) and *Lmx1b^R200Q^*^/+^ (bottom) aggregate footprint profiles subdivided into quintile bins. The range of log2FC changes for each quintile is shown above (*N246K*) or below (*R200Q*) each profile.

We then plotted the magnitude of log2FC footprint score changes for all bound Lmx1b motifs and found, unexpectedly, a continuum of losses, gains, or unchanged scores in *Lmx1b^N246K/+^* and *Lmx1b^R200Q/+^ Pet1* neuron epigenomes (Fig. 3B, C). Footprint score changes in the missense heterozygotes resulted from combined changes in footprint protection and flanking accessibility. Although the overall profiles of *Lmx1b^N246K^* and *Lmx1b^R200Q^* heterozygosity on Lmx1b motif binding activity and flanking accessibility were similar in the two missense mutants, N246K forced a greater proportion (59%) of decreased footprint scores than R200Q, while R200Q forced a greater proportion (57%) of gains (Fig. 3B, C). In *Lmx1b^N246K/+^*, 1433 footprints had fold changes more negative than −1.5X compared to 1184 footprints with this magnitude change in *Lmx1b^R200Q/+^*.

To further illustrate the range of footprint score changes, we binned by magnitude and direction of change, the log2FC into quintile aggregate plots (Fig. 3D, E). Lower Quintile 2 (LQ2) displays Lmx1b motif aggregate footprints that were completely lost, thus unbound, in missense heterozygotes while LQ1 shows milder aggregate footprint losses. The middle quintile (MQ) aggregate plot illustrates motifs in which footprint scores changed very little or not at all. Motifs in Upper Quintile 1 (UQ1) showed either no changes or mild increases in the footprint scores. Indeed, there were more than 1000 bound Lmx1b motifs between MQ and UQ1 that showed no change in footprint score in either heterozygous mutant. UQ2 displays Lmx1b motif footprint scores that were increased in the missense heterozygotes resulting from increases in flanking accessibility and footprint protection. UQ2 also includes 1101 new Lmx1b motif footprints in *Lmx1b^N246K/+^* and 1194 new footprints in *Lmx1b^R200Q/+^*that were not observed in control *Pet1* neuron chromatin; these had a mean footprint score increase of 0.88X log2FC for *Lmx1b^N246K/+^* and 1.24X log2FC for *Lmx1b^R200Q/+^*. These findings indicate that missense heterozygosity forced a complex combination of complete elimination, partial losses, no change, or gains in Lmx1b motif binding activity and flanking chromatin accessibility.

The foregoing results prompted us to investigate whether the DNA binding null allele, Δ, would also force complex losses and gains on Lmx1b motif footprints. Similar to *Lmx1b^N246K/+^* and *Lmx1b^R200Q/+^*, the *Lmx1b*^Δ*/+*^ aggregate footprint profile was decreased compared to the control footprint profile (Fig. S10A). Qualitatively, we found when plotting *Lmx1b*^Δ*/+*^ log2FC footprint score changes, a similar profile of losses, gains, or unchanged Lmx1b motif footprint changes. However, despite lacking the entire HD, Δ was not as potent as the third helix point mutant, N246K, in forcing footprint losses and was similar to R200Q in forcing a greater proportion of gains versus losses (Fig. S10B). Consistent with these DGF results, we found that similar to *Lmx1b^R200Q/+^*, the number of down DEGs in *Lmx1b*^Δ*/+*^ *Pet1* neurons was substantially less than in *Lmx1b^N246K/+^ Pet1* neurons and *Lmx1b*^Δ*/+*^ up DEGs outnumbered *Lmx1b*^Δ*/+*^ down DEGs (Fig. S10C, D). Similar to the missense mutants, GO analysis indicated that *Lmx1b*^Δ*/+*^ DEGs were enriched for synapse genes (Fig. S10E). The smaller total number of DEGs may have hindered enrichment for additional SAM-related terms because when manually annotated we found that 34% of *Lmx1b*^Δ*/+*^ DEGs were associated with SAM-related terms. Unlike the missense mutant DEGs, *Lmx1b*^Δ*/+*^ DEGs were enriched for both chemical synaptic transmission and neurotransmitter transport terms resulting from decreased expression of 5-HT transporter genes, *Slc6a4 and Slc22a3* (Fig. S10E, F), suggesting potentially greater loss of 5-HT reuptake capacity with the Δ allele (Fig. S10F). Despite the lower number of *Lmx1b*^Δ*/+*^ DEGs, we found 61 common DEGs, of which 33% were SAM genes, in *Lmx1b*^Δ*/+*^ vs *Lmx1b^N246K/+^* transcriptomes (Fig. S10G), which may account for their similar axon arbor phenotypes.

The total elimination of some Lmx1b footprints together with substantial gains in missense and Δ heterozygous *Pet1* neurons was surprising given the kinds of mutations we introduced into the Lmx1b HD. In light of this counterintuitive result, we wondered whether the impact of missense and Δ mutants on the *Pet1* neuron epigenome results from combined loss of Lmx1b motif binding together with mutant interference of remaining wildtype Lmx1b function. To investigate this idea, we performed DGF on *Lmx1b^f/f;Pet1-Cre;Ai9^* ATAC-seq datasets (14) in which sequence specific Lmx1b binding is completely eliminated thus mimicking the condition of maximal mutant protein interference of wildtype protein. Contrary to our a priori assumption that Lmx1b motif footprints would be uniformly severely reduced or eliminated in *Lmx1b^f/f;Pet1-Cre;Ai9^ Pet1* neuron chromatin, we found a continuum of footprint losses and gains (Fig. S10H). Notably, the proportion of losses was greater for *Lmx1b^f/f;Pet1-Cre;Ai9^* than found in *Lmx1b^N246K/+^* as expected given the homozygous loss of Lmx1b motif binding. This result supports the idea that missense Lmx1b proteins interfere with wildtype Lmx1b resulting in footprint losses greater than expected from simple heterozygous DNA binding dosage reduction. Further, even homozygous loss of Lmx1b motif binding forced footprint gains perhaps by creating an anomalous epigenomic condition in which other expressed HD TFs gain access to some Lmx1b motifs or nearby flanking motifs resulting in increased TF binding activity. Lmx1b is the only LIM HD expressed in *Pet1* neurons (13, 15, 27, 42). However, several other HD TFs are expressed in *Pet1* neurons, and some are up DEGs in both missense heterozygotes. Hence, we performed a correlation analysis of all HD motifs to the Lmx1b motif. A total of 39 HD TF motifs were highly similar to the Lmx1b motif including *Pou3f2*, which is an up DEG in *Lmx1b*^Δ*/+*^ and *Lmx1b^R200Q/+^ Pet1* neurons (Pearson correlation coefficient > 0.6, p-value < 0.01) (Fig. S10I). This supports the possibility that other *Pet1* neuron expressed HD TFs bind to UQ2 Lmx1b motifs in heterozygous missense *Pet1* neurons.

These results suggest that the missense and Δ proteins are stably expressed in 5-HT neurons. To investigate this, we performed immunostaining of midbrain tissue sections with a polyclonal Lmx1b antisera that specifically co-stained virtually all 5-HT positive neurons in the dorsal raphe and TH^+^ neurons in the SNc (Fig. S11). Despite complete loss of 5-HT immunostaining (Fig. S3A), robust Lmx1b antibody staining was detected in the two ventral longitudinal domains coinciding with the location of developing *Pet1* (5-HT) neurons in *Lmx1b^N246K^*^/*N246K*^, *Lmx1b^R200Q/R200Q^* and *Lmx1b*^Δ*/*Δ^ embryos (Fig. S11). The reduced numbers of Lmx1b^+^ cells in *Lmx1b^N246K^*^/*N246K*^ and *Lmx1b*^Δ*/*Δ^ compared to control and *Lmx1b^R200Q/R200Q^*is consistent with the reduced numbers of TdTomato^+^ *Pet1* neurons in *Lmx1b^N246K^*^/*N246K*^ and *Lmx1b*^Δ*/*Δ^ embryos (Fig. S2B, D).

We next sought to identify distinguishing features that may account for the dramatically different changes in Lmx1b motif binding activity and flanking accessibility resulting from missense heterozygosity. We found that the magnitude and direction of footprint score changes in both missense heterozygotes correlated with the level of Lmx1b motif binding activity in control *Pet1* neurons, with LQ2 motifs showing the highest level of binding activity and UQ2 the lowest (Fig. 4A, B). Lmx1b CUT&RUN data revealed a similar difference in Lmx1b occupancy among the five groups with LQ2 showing the greatest level of occupancy and UQ2 the lowest (Fig. 4C, D). In addition, a greater number of Lmx1b motifs with footprints in LQ2 showed CUT&RUN peaks (31%, 248/790) compared to UQ2 Lmx1b motifs with footprints (7.3%, 58/790). Given the higher Lmx1b occupancy in LQ2 vs UQ2, we wondered whether there was an enrichment for additional Lmx1b motifs in LQ2 accessible chromatin. We first masked central Lmx1b motifs in footprints, then searched the flanking ±250 bp with a set of HD motifs (43). We found that LQ2 Lmx1b footprints were highly enriched for additional Lmx1b and other HD motifs in flanking accessible sequences compared to scrambled background sequences in both missense heterozygous *Pet1* neurons (Fig. 4E, F). In contrast, UQ2 Lmx1b footprints did not show an enrichment for Lmx1b motifs in flanking sequences and displayed a much weaker enrichment for other HD motifs (Fig. 4G, H). Since flanking regions of LQ2 footprints are enriched for Lmx1b and other HD TF motifs, we next asked whether LQ2 Lmx1b motif footprints were more likely to be located next to other LQ2 Lmx1b motif footprints in the same CREs. Indeed, there were more than 2X (*N246K/+*) and 1.46X (*R200Q/+*) as many LQ2 Lmx1b motif footprints less than 500bp away from another LQ2 Lmx1b motif footprint compared to footprints in the other quintiles (Fig. 4I). Thus, Lmx1b motifs featuring the strongest level of motif binding activity, Lmx1b occupancy, and enrichment for flanking Lmx1b motifs and footprints in control *Pet1* neurons are those that exhibit complete loss of Lmx1b binding activity and flanking accessibility in missense heterozygotes.

**Figure 4:**
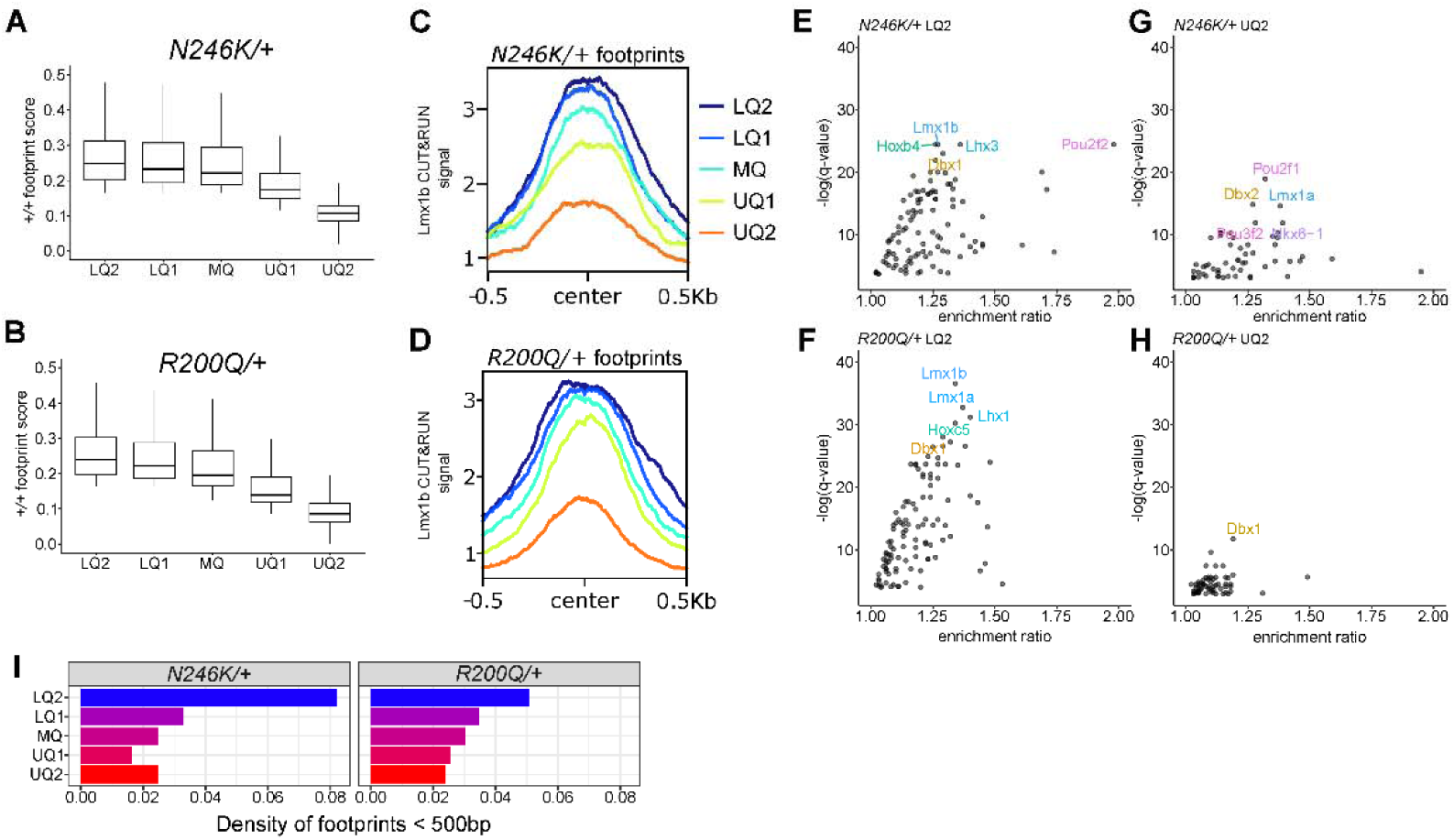
Distinguishing features of lost and gained Lmx1b footprints. A-B) Lmx1b motif binding activity (footprint scores) in control *Pet1* neurons according to missense, *Lmx1b^N246K/+^* (A) and *Lmx1b^R200Q^*^/+^ (B) motif quintiles. UQ2 motifs show the least Lmx1b motif binding activity in control *Pet1* neurons compared to motifs in other quintiles. C-D) Lmx1b CUT&RUN signal at E14.5 in control *Pet1* neurons at Lmx1b motifs segregated by quintile, *Lmx1b^N246K/+^* (C) and *Lmx1b^R200Q^*^/+^ (D). Lmx1b occupancy in control neurons is greatest at LQ2 motifs. E-H) Enrichment of Lmx1b and other HD motif adjacent to LQ2 Lmx1b motifs in *Lmx1b^N246K/+^* (E) and *Lmx1b^R200Q^*^/+^ (F) and adjacent to UQ2 Lmx1b motifs in *Lmx1b^N246K/+^* (G) and *Lmx1b^R200Q^*^/+^ (H). There is a greater level of Lmx1b and HD motif enrichment at LQ2 Lmx1b motifs compared to UQ2 Lmx1b motifs. I) Density of Lmx1b motif footprints in each motif quintile for *Lmx1b^N246K/+^* (left) and *Lmx1b^R200Q^*^/+^ (right) within 500bp of another footprint.

We previously obtained ChIPmentation data for H3K4me3 (promoter), H3K4me1 (enhancer), H3K27ac (activation), and H3K27me3 (repression) marks from E14.5 *Pet1* neurons to define chromatin states across the *Pet1* neuron genome (14). Here, we used these datasets to determine whether the magnitude and direction of Lmx1b footprint changes in missense heterozygotes were associated with particular chromatin states. We found a greater enrichment for active, TSS-distal enhancer and active promoter signatures in LQ2 footprints in both missense mutants compared to other groupings, with UQ2 footprints showing the least enrichment (Fig. S12A). In contrast, UQ2 footprints showed a greater enrichment in the other/no signal chromatin state signature (Fig. S12A). These data suggest that Lmx1b footprints that are completely eliminated by missense heterozygosity are more frequently located in active, TSS-distal enhancer and active promoter regions, while Lmx1b footprints exhibiting the greatest gains in footprint scores are more frequently located in chromatin lacking active cis-regulatory histone mark signatures.

To identify potential functions of genes linked to missense altered Lmx1b footprints, we associated each footprint to the nearest gene TSS within ±50kb. We then performed GO term enrichment analysis to determine potential functions of footprint linked genes within each quintile. This analysis did not reveal dramatic differences in GO term enrichment among the quintiles. However, we found that the top significantly enriched terms in each quintile robustly converged on processes related to synapse and axon development (e.g. synapse organization, neuron projection morphogenesis, cell projection morphogenesis) (Fig. S12B). Further, we identified a range of nearby Lmx1b motif footprint score changes associated with SAM gene DEGs, whose expression was commonly significantly altered in both missense heterozygotes (Fig. S12C). These findings suggest that altered Lmx1b binding activity at CREs results in altered SAM gene expression.

### Lmx1b missense disruption of *Pet1* neuron GRNs

We investigated the impact of Lmx1b missense on Lmx1b-dependent TF networks in *Pet1* neurons. Differential expression analysis of RNA-seq datasets identified many common significantly upregulated TFs in *Lmx1b^N246K^*^/+^ and *Lmx1b^R200Q^*^/+^ mutants, suggesting that heterozygous missense mutations force derepression of different *Pet1* neuron TF networks. DGF revealed changes in Lmx1b motif binding activity within ± 50kb of the TSSs of these derepressed TF genes. A broad range of Lmx1b footprint changes encompassing LQ and UQ footprint quintiles were found associated with the upregulated TFs (Fig. 5A). Most of the footprint changes occurred in TF gene introns or within 2kb of the TF’s TSS. For example, the substantial upregulation of *Sox21* was associated with a complete footprint loss (LQ2) in the immediate 3’ flanking region of *Sox21* suggesting missense interference of Lmx1b motif binding activity at a CRE that may normally repress transcription.

**Figure 5:**
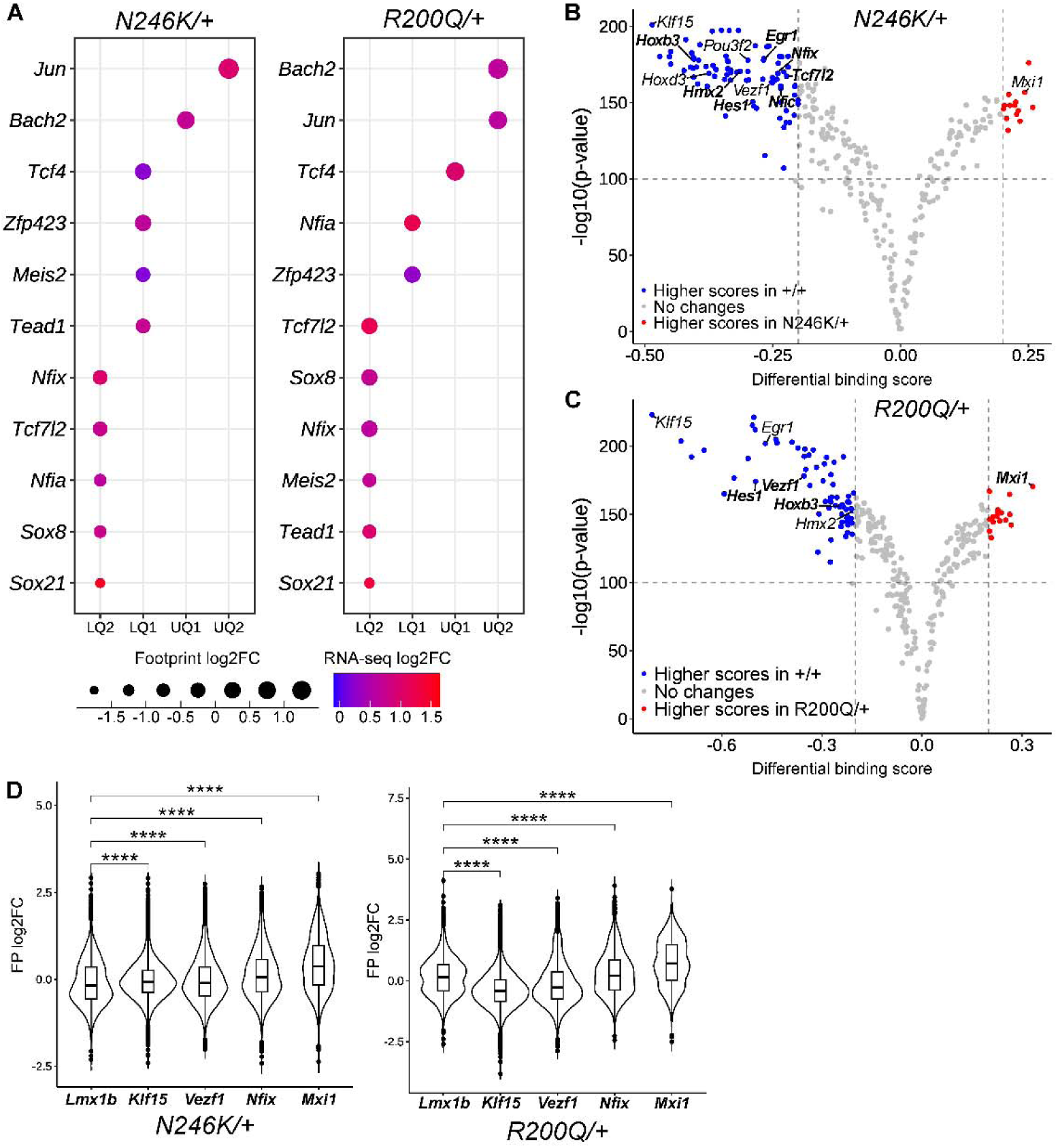
L*m*x1b heterozygous missense mutations disrupt Lmx1b-dependent GRNs in *Pet1* neurons. A) Dot colors represents different magnitudes of individual TF DEG up regulation in the indicated heterozygous genotype according to the log2FC expression heatmap. Dot size represents the log2FC magnitude and direction of footprint change found associated with the indicated TF DEG sorted into the appropriate footprint change quintile. B-C) Altered TF motif binding activities in *Lmx1b^N246K/+^* (B) and *Lmx1b^R200Q^*^/+^ (C). Each dot represents an aggregate footprint score change for a single TF binding motif; to enhance biological relevance analysis was limited to motifs whose associated TF is expressed at > 1 FPKM in *Pet1* neurons. Blue dots indicate decreased TF motif binding activity in missense heterozygous *Pet1* neurons vs. activity in control *Pet1* neurons (+/+). Red dots indicate increased TF motif binding activity in missense heterozygous *Pet1* neurons vs. activity in control *Pet1* neurons (+/+). Bold lettering indicates TF DEGs in the indicated heterozygous genotype; non-bold lettering indicates TF gene expression was not found altered in the indicated heterozygous genotype. Differential binding score: aggregate log2FC for each motif. D) Violin plots of individual log2FC footprint bound motif changes for the indicated TF in *Lmx1b^N246K/+^* (left) and *Lmx1b^R200Q/+^* (right) *Pet1* neuron chromatin.

We next investigated whether heterozygous Lmx1b missense mutations disrupted motif binding activity of other *Pet1* neuron expressed TFs. We found decreased aggregate footprint scores for 81 different TF motifs while increased aggregate scores were found for 14 different TF motifs in *Lmx1b^N246K^* heterozygotes (Fig. 5B). In *Lmx1b^R200Q^* heterozygotes, decreased and increased motif binding activity was found for 66 and 16 expressed TFs, respectively (Fig. 5C). Footprint score changes in both missense mutants resulted from changes in motif protection and flanking accessibility. Plotting individual log2FC footprint score changes across all bound motifs revealed unique profiles of losses and gains for each expressed TF in *Lmx1b^N246K/+^* and *Lmx1b^R200Q/+^ Pet1* neuron epigenomes (Fig. 5D). For example, decreased footprint scores predominated for Klf15 motif binding activity, particularly in *Lmx1b^R200Q^* heterozygotes. In contrast, increased footprint scores predominated for Mxi1 motif binding activity in both missense mutants (Fig. 5D). Together, these findings indicate that *Lmx1b^N246K^* and *Lmx1b^R200Q^* heterozygous missense mutations uniquely disrupt not only Lmx1b motif binding activity, but also the binding activity of various expressed TFs that likely compose *Pet1* neuron GRNs.

## Discussion

Here, we present mouse lines that model two different recurrent pathogenic LMX1B HD missense mutations; we sought to understand the impact of LMX1B missense mutations on 5-HT neuron development. Homozygous germ line and conditional targeting of *Lmx1b* has revealed its multiple key embryonic and early postnatal stage regulatory roles in the development of mouse 5-HT neurons (7, 13, 44). It is, therefore, noteworthy that missense heterozygosity had no impact on any of the several regulatory steps we examined that Lmx1b controls during embryonic 5-HT neurons development. *Lmx1b^N246K^* and *Lmx1b^R200Q^* heterozygosity had no effect on development of *Pet1* neuron precursor numbers, 5-HT primary axon growth and secondary axon projections to distal target region of the forebrain. 5-HT^+^ and Tph2^+^ neuron numbers also did not differ from control numbers in any of the raphe nuclei, although the moderately reduced levels of *Tph2* and *Ddc* expression, particularly in *Lmx1b^N246K^*, may result in quantitatively reduced 5-HT synthesis. Further, missense heterozygosity did not disrupt development of TH^+^ DA neurons in the SNc or TH fibers in the striatum. These findings indicate that a single wildtype Lmx1b copy appears sufficient for execution of many Lmx1b-dependent roles in neuron development.

In contrast, we found a forebrain-wide deficit in 5-HT axon arbors, most notably in the motor cortex and hippocampus of both missense heterozygous mice. The axon deficits in the missense heterozygotes occurred in the P7-P30 period and therefore resulted from failed formation of arbors and not formation of major serotonergic projection pathways or post-developmental arbor degeneration. Despite the massive deficit in 5-HT axon arbors in the spinal cord in homozygous *Lmx1b* targeted mice (13), axon densities in the missense heterozygous spinal cord were not statistically different from control revealing yet another Lmx1b-dependent developmental step that appears insensitive to Lmx1b dosage. RNA-seq of heterozygous missense *Pet1* neuron transcriptomes supported a requirement for two intact copies of *Lmx1b* in the development of *Pet1* neuron transcriptomes, particularly for expression of synapse, axon, and mitochondrial related genes. The greater number of *Lmx1b^N246K/+^* DEGs and the large number not shared with *Lmx1b^R200Q/+^* DEGs may account for the SVZ and striatum axon arbor densities differences in two the lines.

What might explain the highly selective sensitivity of postnatal forebrain axon arbor formation to Lmx1b missense heterozygosity? One possibility is that the weeks long stage of early postnatal 5-HT axon arborization, which generates an extraordinary increase in total single neuron axon length, places a greater demand on Lmx1b output for SAM gene expression to build the profuse terminal extensions of the neuron (45, 46). Thus, while Lmx1b transcriptional output in missense heterozygotes is largely sufficient at embryonic stages, it may be inadequate for expansive postnatal forebrain arborization. An alternative non-mutually exclusive possibility is that Lmx1b transcriptional activity is downregulated at postnatal stages consequently rendering single allele *Lmx1b* transcriptional output below threshold for adequate SAM transcription (47).

Biallelic stop gain mutations that destroy the DNA binding capacity of FEV (human Pet1 ortholog) were identified in brothers with autism spectrum disorder and severe intellectual disability suggesting disruption of human serotonergic GRNs may contribute to neurodevelopmental disorders (48). Similarly, our findings suggest that human heterozygous LMX1B missense mutations may disrupt the regulatory program that controls human 5-HT neuron development. Humans appear highly sensitive to LMX1B dosage as heterozygous missense individuals have skeletal, kidney, and ocular pathology. Given this dosage sensitivity, it seems reasonable to predict axon arbor density alterations in at least some of these individuals. However, this would be much more difficult to determine than the easily detectable skeletal, kidney, and ocular pathology. Individuals with heterozygous LMX1B mutations were reported to show increased symptoms of attention deficit hyperactivity disorder and major depressive disorder (49). We found a disruption of spatial memory in *Lmx1b^R200Q^* heterozygous females, which was associated with reduced 5-HT axon arbor densities in the SLM, previously implicated in modulating spatial memory formation (34). These findings suggest relatively modest developmental deficits in postnatal 5-HT arborization can result in cognitive impairments. The well-established intra- and interfamily heterogeneity in peripheral disease severity and penetrance in human LMX1B cases raises the possibility that human missense heterozygosity could be associated with stress-related behavioral phenotypes in the absence of peripheral disease. Further neuropsychiatric evaluation of heterozygous LMX1B individuals may be informative. Given 5-HT’s broad influence on circuitry underlying cognitive and emotion-related behaviors perhaps human 5-HT axon arbor deficits impact temperament or personality traits that are less readily modeled or assayed in rodents (50, 51).

Application of DGF in developing in heterozygous missense *Pet1* neurons provided unexpected insight into the poorly understood question of how stably expressed TF missense proteins impact the epigenome of a specific neuron-type, in vivo. We found that missense heterozygosity forced a continuum of losses and gains in Lmx1b footprint magnitudes. Missense heterozygosity completely eliminated footprints and flanking accessibility at about one quarter of bound Lmx1b motifs. These lost Lmx1b motif footprints were distinguished from other missense altered Lmx1b footprints by exhibiting the strongest level of Lmx1b motif binding activity in control *Pet1* neurons, the strongest level of Lmx1b occupancy, enrichment for additional Lmx1b motifs and footprints in flanking accessible sequences, and greater enrichment for active distal enhancer and promoter signatures. Missense heterozygosity also produced partial losses or no change in footprints at other sets of motifs. Finally, missense heterozygosity caused increased footprints at a large set of distinct Lmx1b motifs. Motifs with the strongest gains featured the weakest Lmx1b motif binding activity in control *Pet1* neurons, the lowest level of Lmx1b occupancy and no enrichment for additional Lmx1b motifs in flanking accessible sequences. Despite the varied types of changes, each group of altered footprints were associated with genes that strongly converged on synapse and axon processes, further corroborating missense disruption of axon development.

DGF further revealed that Lmx1b missense mutations broadly disrupted Lmx1b-dependent TF networks in *Pet1* neurons. We found a spectrum of footprint losses and gains at motifs for disparate TFs expressed in *Pet1* neurons. Expression of some of these TFs was altered in the missense heterozygotes suggesting footprint changes result directly from altered TF expression. The expression levels of other TFs were not found to be changed in the missense heterozygotes suggesting footprint changes may result from expression changes in other epigenetic regulators that are needed for motif accessibility. The identification of which footprint changes are directly responsible for gene expression changes and arbor defects will require perturbation of different TF motifs most likely requiring combined motif targeting in the face of possible CRE redundancy.

The heterozygous missense axon and transcriptome phenotypes give the appearance of haploinsufficiency when compared to the much more severe axon and transcriptome phenotypes evident in homozygous conditionally targeted of *Lmx1b* mice (13). This would typically be interpreted as simple dosage reduction in gene product resulting in a corresponding uniform reduction in function. However, our DGF results suggest a different mode of action. Given the kinds of point mutations we introduced into the HD third (recognition) helix or the HD N-terminal arm, it is likely that they create mutant proteins that are null (N246K) or near null (R200Q) loss of DNA binding to Lmx1b motifs; the multiple anatomical similarities we report, particularly for *Lmx1b*^Δ*/+*^ and *Lmx1b^N246K/+^*, is corroborative. However, DGF strongly suggests that the seeming axon and transcriptome haploinsufficiency phenotypes do not result simply from elimination of DNA binding in one Lmx1b copy in view of the large number of Lmx1b motif footprints that were completely lost in mutant heterozygotes. LIM HD function depends on complex formation with cofactors such as the multi-adaptor LIM domain binding proteins (LDB) or other sequence-specific TFs (52, 53). Hence, we suggest that expressed missense proteins not only cannot bind Lmx1b motifs, but they also interfere with remaining wildtype Lmx1b protein in forming functional complexes with cofactors (Fig. S13). DGF analysis of *Lmx1b^f/f;Pet1-Cre;Ai9^*, in which Lmx1b motif binding is completely lost and therefore mimics maximal missense interference of wildtype provides support for “antimorphic” action of heterozygous Lmx1b missense. The more severe axon and transcriptome phenotypes and greater proportion of footprint losses in *Lmx1b^f/f;Pet1-Cre;Ai9^* suggest heterozygous missense interference is not maximal and some functional Lmx1b complexes remain. DGF further revealed an additional effect of missense heterozygosity, not previously associated with haploinsufficiency: secondary gains in TF motif binding activity. The increase in Lmx1b motif footprints at some bound Lmx1b motifs in heterozygous missense, Δ, and even *Lmx1b^f/f;Pet1-Cre;Ai9^ Pet1* neurons suggests that loss of Lmx1b binding enables other TFs (expressed HDs and non-HDs) to bind at some Lmx1b motifs and/or at flanking accessible motifs resulting in footprint gains. The greater number of footprint losses in *Lmx1b^N246K/+^* compared to *Lmx1b^R200Q/+^*and *Lmx1b*^Δ*/+*^ losses and the fewer number of gains in *Lmx1b^N246K/+^* suggest N246K exerts stronger interference of wildtype protein and/or other TFs whose binding may be enabled in the mutant *Pet1* neurons. Thus, DGF suggests the outward appearance of haploinsufficiency in missense heterozygotes results from complex losses and gains of TF motif binding activity in *Pet1* neuron epigenomes.

## Materials and Methods

The detailed experimental procedures including mouse lines, histology, quantification of axon densities, cell body counts, behavior, cell sorting, INTACT, RNA-seq, ATAC-seq, DGF, GO analysis, ChIPmentation, and statistical analysis are comprehensively described in SI Appendix, Materials and Methods.

## Acknowledgments

Funding: R01MH125918, R01MH117643; K01NS135573 to N.T. This work was supported by the Cytometry and Microscopy Shared Resource at Case Comprehensive Cancer Center, the CWRU Genomics Core, and the CWRU Mouse Behavior Phenotyping Core.

