## Supporting information appendix for "*LMX1B* missense-perturbation of regulatory element footprints disrupts serotonergic forebrain axon arborization"

Evan S. Deneris

William C. Spencer

**Materials and Methods**

**Mouse lines**

Mice were maintained according with the National Institutes of Health Guidelines for the Care and Use of Laboratory animals and were approved by the CWRU Institutional Animal Care and Use Committee. Experiments were performed with male and female mice using age-matched controls. Littermate controls were used whenever possible.

*Missense mouse line generation*: CRISPR/Cas9 genome editing was performed in the CWRU Transgenic and Targeting Facility. Subsequent to the description of the original mutations identified in *LMX1B*, an additional 69bp was identified at the 5’ end of first coding exon of the human gene. This results in a 23 amino acid difference in human and mouse Lmx1b primary structure numbering. Therefore, the original p.R200Q and p.N246K human mutations are corrected to p.R223Q ([NM_001174147.2:c.668G>A](https://www.ncbi.nlm.nih.gov/nuccore/NM_001174147.2?report=graph&search=NM_001174147.2%3Ac.668G%3EA) ) and p.N269K ([NM_001174147.2:c.807C>A](https://www.ncbi.nlm.nih.gov/nuccore/NM_001174147.2?report=graph&search=NM_001174147.2%3Ac.807C%3EA)). The additional 69bp human sequence is not present in the mouse genome, therefore the orthologous mouse mutations are designated as p.R200Q and p.N246K. To generate human mutation knock-in mice, sgRNAs targeting mouse *Lmx1b* exons 4 and 5, were designed using CRISPOR selecting sgRNAs with the lowest number of possible off-target effects (1). sgRNA for *Lmx1b^N246K^*: 5’- GACCTCTCACCTTTGCTCTT-3’, sgRNA for *Lmx1b^R200Q^*: 5’-GTCTCATCGCTGCCGGGGAC-3’. sgRNAs and Cas9 from *S. pyogenes* with an NLS were ordered from PNA Bio. Templates for homology-directed repair were designed with 50 base regions of homology flanking the targeted nucleotide and were ordered as PAGE purified Ultramers from IDT. The PAM sites were mutated preventing recutting.

*Lmx1b^N246K^* template: 5’CAGCAGAGACAGGCCTCAGCGTGCGTGTGGTCCAGGTCTGGTTTCAG**AAA**CAAAGAGCAAAGGTGAGAGGTCACGACTAATCCAGTGCTCCAGGGCAGAC-3’.

*Lmx1b^R200Q^* template: 5’TAAAGGCAGTGGAGATGACGGGAAAGACCCGAGAAGGCCCAAACGGCCT**CAA**ACCATCCTCACCACACAGCAGCGAAGAGCTTTCAAGGCATCCTTTGAG-3’.

The Cas9 protein, sgRNA, and HDR template oligo were mixed at 10ng/µl each and injected into fertilized eggs from B6.SJL mice. A custom Taqman genotyping assay was designed to identify each nucleotide change (ThermoFisher). Founders were crossed with C57Bl/6J mice to determine the animals showing germline transmission of point mutations. Exons 4, 5, and 6 of *Lmx1b* containing the homeodomain were PCR-amplified and Sanger sequenced to confirm there were no other mutations except the intended AA**C** to AA**A** for *Lmx1b^N246K^* and C**G**A to C**A**A for *Lmx1b^R200Q^*. Mice were backcrossed with C57Bl/6J mice to establish the lines.

*Lmx1b*^Δ/Δ^ mice were generated by using the *Dbx1-Cre* (2) and *Lmx1b^fl^* (3) mice to induce germline recombination (4) of *Lmx1b* exons 4-6, thereby removing the entire homeodomain and some flanking amino acids. *Dbx1-Cre* was removed in subsequent generations. Conditional fluorescent labeling was achieved using the *Pet1-Cre* (5) and *Ai9* (*Rosa*^Tom^; Jackson lab) alleles.

**Histology and Immunofluorescence**

Mice (P7, P30, adult) were anesthetized with Avertin (44 mM tribromoethanol, 2.5% tert-amyl alcohol, 0.02 ml/g body weight) and transcardially perfused with ice cold phosphate-buffered saline (PBS) for 2 min, followed by 4% paraformaldehyde (PFA) in PBS for 15 min.  Brains and spinal cords were removed and post-fixed in cold 4% PFA in PBS for 2 hr, followed by 30% sucrose/PBS overnight at 4°C. E18.5 embryos were transcardially perfused with 4 ml of 4% PFA and post-fixed in 4% PFA overnight at 4°C, followed by 30% sucrose/PBS overnight at 4°C. E13.5 embryos were drop-fixed in 4% PFA overnight at 4°C, followed by 30% sucrose/PBS overnight at 4°C. Tissue was embedded in Optimal Cutting Temperature compound, flash frozen in liquid nitrogen, and sectioned on a cryostat at 25 µm. Tissue sections were mounted on MAS-GP Adhesive Glass Slides and vacuum dried. Sections were then permeabilized in 0.3% Triton 100X-PBS (PBS-T) for 15 min followed by antigen retrieval in sodium citrate buffer for 5 min in a microwave oven at low power. Sections were blocked with 10% Normal Goat Serum in PBS-T for 1 hr followed by incubation in primary antibody at 4°C overnight. Secondary antibodies were used for 1 hr at room temperature.  Primary antibodies used were mouse or rabbit anti-RFP, rabbit anti-Tph2, rabbit anti-5-HT, guinea anti-Lmx1b, and mouse anti-TH. Secondary antibodies used were goat anti-rabbit, mouse, or guinea Alexa Fluor 488, 594, and 647. Sections were treated with DAPI for 5 min at room temperature to visualize nuclei.

**Microscopy**

Immunofluorescent images were captured with an LSM800 confocal microscope or Axio Imager M2 microscope (Carl Zeiss). Brightness and contrast were modified equally between genotypes used for analysis.

**Quantification of Axon Densities**

TdTomato^+^ and TH^+^ axon densities were compared using 2-3 25µm thick sections collected from each animal per genotype. 2-4 month old mice were used for all adult stage analyses. At least 3 biological replicates were used for all quantified experiments, as indicated in the figure legends. For each section, non-overlapping images from comparable regions of the brain were captured with a Zeiss LSM800 with 20x or 40x magnification and stitched to uniform size using Zeiss 2.3 software. ImageJ-Fiji was used to trace regions of interest and measure respective axon densities. Axon densities were normalized to control values that were mounted on the same glass slide.

**Cell Body Counts**

TdTomato^+^ cells in adult (2-4 months) heterozygous mutants were counted in every 4th section taken across the entire rostro-caudal extent of either the DRN, MRN, B9, and medullary raphe. Tph2^+^ and 5-HT ^+^ cells in adult (2-4 months) heterozygous mutants were counted in every 4th section taken across the entire rostro-caudal extent of the DRN. TH^+^ cells in adult (2-4 months) heterozygous mutants were counted in every 4th section across the SNc. TdTomato^+^ cells in E15.5 homozygous mutants were counted in every 2nd section. 3-4 biological replicates were used per genotype.

**Photography**

Mouse eyes were photographed using an iPhone 11. P0 brains were imaged using a Leica MZFLIII fluorescence stereomicroscope.

**Behavior Assays**

Male and female mice (6-12 weeks old) were tested separately after being acclimated to the testing room for at least 30 min. Assays were performed during the light phase and experimenters were blinded to genotype. All arenas were thoroughly cleaned with a 70% ethanol solution between experiments. 14-15 mice were used per genotype. *Open Field*: Mice were placed in a 49.5 cm x 49.5 cm arena with 38 cm high walls in a dimly lit room and allowed to freely explore for 10 min. Activity was recorded using Any-maze video tracking software (Stoelting, CO, USA). Inner zones and outer zones were digitally subdivided. Total distance traveled and time spent in the inner zone were measured.  *Elevated Plus Maze:* Mice were placed in a 76 cm x 76 cm cross with a 6.35 cm wide walkway. Two of the arms were enclosed with 19 cm high walls. Mice were allowed to freely explore for 5 min. Activity was recorded using Any-maze video tracking software. Time spent in the open arms was recorded. *Novel Object Recognition:* Mice were habituated to the environment by placing them individually into a 49.5 cm x 49.5 cm arena with two identical objects on opposite sides. Mice were given 10 min to explore the objects before being returned to their home cages. 24 hrs later, mice were returned to the arena with one previously habituated “familiar object” and one new “novel object.” Time spent interacting with each object was recorded, with “interaction” defined as facing the object, 2cm or closer. Discrimination Index (DI) was calculated as follows: DI = Novel object interaction time – Familiar object interaction time/Total exploration time. *Rotarod performance:* each mouse was placed on a Rotarod machine (Rotamex-5, Columbus Instruments) operating at 4rpm and given 1 min to acclimate. The machine was then set to accelerate, increasing .1rpm every second until the animals fell off the rod, recorded as the “Latency to Fall.” Three trials were conducted for each animal. *Marble Burying:* 8 marbles were placed 4 cm apart within a 20 cm x 30 cm cage on top of 5cm of firmly tamped bedding. Mice were then placed within the cage and left alone for 30 min. The number of marbles buried during this time was recorded, with “buried” being defined as at least 2/3rds of the way covered by the bedding.  *Barnes Maze*: Spatial memory performance was investigated using the Barnes Maze as previously described (6). A 91.44 cm diameter circle area was used with twenty 5.08 cm holes along the perimeter. The arena was digitally subdivided into four equal quadrants, and a single escape hatch was placed underneath one of the holes. Briefly, mice were habituated to the testing arena by placing them in a 3,500ml glass beaker for 30 s. The animals were then gently guided to the escape hatch through the beaker and allowed 3 min to independently enter the hole, where they were allowed to stay for 1 min before returning to their home cages. During training, mice were introduced into the maze through an opaque cardboard box and allowed to explore the maze. “Latency to escape” was measured as the time it took each animal to find the escape hatch. The escape hatch was rotated every three trials to avoid intra-maze odor or visual cues. Five training trials were conducted. 48 hrs following the last training day, mice were reintroduced into the arena without the escape hatch for 2 min. The time spent in the “Target Quadrant,” defined as the quarter of the arena which previously housed the escape hatch, was recorded using Any-maze video tracking software.

**Cell Sorting**

The entire rostrocaudal extent of E17.5 *N246K/+; Pet1-Cre; Ai9*, *R200Q/+; Pet1-Cre; Ai9, Lmx1b^Δ/+^*; *Pet1-Cre; Ai9* hindbrains were dissected to capture Tomato+ *Pet1* neurons from all developing raphe nuclei and placed in cold aCSF solution (3.5 mM KCl, 126 mM NaCl, 20 mM NaHCO3, 20 μM dextrose, 1.25 mM NaH_2_PO4, 2 mM CaCl_2_, 2 mM MgCl2, 50 μM AP-V (Tocris), 29 μM DNQX (Sigma), and 100 nM TTX (Abcam)). Dissected tissues were then incubated in bubbling (95% O2, 5% CO2) aCSF containing 1 mg/mL protease from *Streptomyces griseus* (Sigma) for 15 min at room temperature. Following enzymatic digestion, tissues were transferred to 500 μL cold aCSF/10% FBS solution containing 4 μL DNase I (Invitrogen) and slowly triturated with fine fire-polished glass pipettes to generate a single-cell suspension. Cells were then filtered using a 40 μm filter (BD Biosciences) and sorted on a FACS Aria-SORP digital cell sorter with an 85 μm nozzle. Three biological replicates were used per genotype.

**RNA Sequencing**

E17.5 flow-sorted neurons collected into Trizol LS were isolated using chloroform extraction and with RNA Clean and Concentrator-5 kit (Zymo). Total concentration and quality were assessed using Quantifluor RNA system (Promega) and Fragment analyzer (Advanced Analytics). Samples were converted to cDNA and library construction was performed using the Ovation SoLo RNA-Seq Library Preparation Kit for mouse (Nugen Inc). Libraries were sequenced on a NextSeq 550 (Illumina) with single-end sequencing for 75 cycles.

**RNA-seq analysis**

Raw purity filtered RNA-seq reads were aligned to the mouse genome (UCSC mm10) with Hisat2 v2.1.0 (7) with the parameters “penalty of 12 for a noncanonical splice site, maximum and minimum penalty for mismatch of 1,0, and maximum and minimum penalty for soft-clipping of 3,0”. Duplicate reads were removed using the UMI-based tool nudup (Tecan Genomics). Cufflinks v2.2.2 was used for gene expression quantification and differential expression with a fold change threshold of 1.5-fold and FDR threshold of 0.01 (8, 9). Graphs were generated using ggplot2 v3.4.4 (10). The Venn diagram was created using DeepVenn (11).

**Embryonic (E14.5) nuclei isolation and ATAC-seq**

The INTACT protocol (12) was modified for isolation of *Pet1* neuron nuclei. *N246K/+; Pet1-Cre; Ai9*, *R200Q/+; Pet1-Cre; Ai9, Lmx1b^Δ/+^*; *Pet1-Cre; Ai9* mice were crossed with homozygous *R26-CAG-LSL-Sun1-superfolding GFP-Myc* mice (Sun1-GFP). The entire E14.5 hindbrain was microdissected in cold PBS to collect TdTomato^+^ *Pet1* neurons from all developing raphe nuclei. Dissected tissues were Dounce homogenized using a pestle in homogenization buffer. IGEPAL-630 solution was added to bring the homogenate to 0.3% IGEPAL-630. The sample was filtered through a strainer, washed with 1x PBS, and centrifuged at 600g for 5 min. Pellets were mixed with 1 ml HSB buffer, and centrifuged at 10,000g for 10 min. Pellets were incubated with 15 µl Protein A/G Dynabeads (88802, ThermoFisher) and 1 µL rabbit anti-GFP antibody (A-11122, Invitrogen) in 200 µl wash buffer for 1 hr. Beads with GFP^+^ nuclei were washed with 1 ml wash buffer 3 times. Then, beads were resuspended in 25 μL Transposition mix (25 μL 2× TD buffer [Diagenode], 2.5 μL transposase [Diagenode], 22.5 μL H2O) and incubated at 37°C for 25 min on a Thermomixer at 1000 rpm. The resulting DNA fragments were purified using the DNA Clean and Concentrator-5 Kit (Zymo) and PCR amplified with Illumina Nextera adapter primers using the NEBNext High Fidelity 2× Master Mix (NEB). Final PCR products were cleaned using PCRClean Dx beads (Aline Biosciences), assessed for quality on a Bioanalyzer, and sequenced on an Illumina NextSeq 550. Two replicates were generated for *Lmx1b^+/+^*, *Lmx1b^R200Q/+^* and *Lmx1b^Δ/^* and one replicate for *Lmx1b^N246K/+^*.

**ATAC-seq data processing**

ATAC-seq reads were mapped and filtered using the standardized ENCODE consortium ATAC-seq pipeline v2.2.2 (13). Purity filtered sequencing reads were mapped to the mouse genome (UCSC mm10) with Bowtie2 v2.3.4.3. Duplicate reads were marked with Picard v2.20.7 and removed with Samtools v1.9. Mitochondrial reads were mapped with Bowtie2 as above and removed with Samtools. Biological replicate read data for *Lmx1b^N246K/+^*, *Lmx1b^R200Q^*^/+^ and *Lmx1b^Δ/+^* were merged with Samtools and control, *Lmx1b^N246K/+^*, *Lmx1b^R200Q^*^/+^ and *Lmx1b^Δ/+^* mapped reads were downsampled to 100M reads each with sambamba v1.0 (14).

**Digital genomic footprinting analysis**

ATAC-seq footprinting analysis was performed with TOBIAS v.0.16.0 (15). Peaks of accessible chromatin regions for each sample were called with MACS2 using the parameters “--gsize mm --nomodel --shift -100 --extsize 200 -broad”. Peaks were annotated to genes using UROPA v4.0.2 (16) up to a limit of 50 kb. Since the Tn5 transposase has an inherent bias in DNA sequence cutting preference, expected Tn5 cuts were subtracted from the uncorrected ATAC-seq signal. Motifs from the JASPAR 2022 CORE non-redundant vertebrate collection (17) were mapped to the genome and footprinting scores were calculated for each motif that are located with accessible peak regions. Differential binding/activity scores for each motif were calculated between each control, *Lmx1b^N246K/+^*, *Lmx1b^R200Q^*^/+^, *Lmx1b^Δ/+^* and *Lmx1b^f/f;Pet1-Cre;Ai9^* samples and filtered for TFs that were detected as expressed in the RNA-seq data (>1 FPKM). Aggregate footprint plots were generated with TOBIAS visualization modules. Heatmap and volcano plots were generated using ggplot2 v3.4.4 in R v4.3.2 (10).

**Annotation of footprints to genes**

Footprints were annotated to genes using a proximal region of 5kb upstream and 1kb downstream for each gene and extending to the proximal neighboring gene region or up to 50kb in one direction using GREAT (18, 19).

**Gene ontology enrichment**

Gene ontology enrichment analysis was performed with Webgestalt with at least five genes per category and FDR <= 5% (20). The enriched GO term bubble plots were graphed using ggplot2 v3.4.4 in R v4.3.2. The GO term gene count was performed in R v4.3.2 and graphed using ggplot2 v3.4.4 (10). We used all three categories: Biological process, Molecular function, and Cellular Component.

**Footprint density analysis**

The distance from each footprint to the next neighboring footprint was calculated using the GenomicDistributions package v1.9.0 in R v4.3.2 (21). The distances for footprints in each quintile were filtered for footprints that were within 500bp of the neighboring footprint. The density of footprints for each quintile was plotted as the count of footprint distances <500bp divided by the total number of footprints in each quintile using R using ggplot2.

**ChIP-seq using Tagmentation (ChIPmentation)**

*Pet1* neurons were flow sorted from *Pet1-EYFP* embryos at E14.5 and ChIPmentation was performed as previously described (22). In brief, sonicated chromatin was incubated with anti-CTCF antibody on a rotator overnight at 4°C. Antibody bound chromatin was washed then Protein A/G dynabeads were added and incubated for 2 hr at 4°C on a rotator. Beads were washed then resuspended in tagmentation mix containing tagmentation buffer and Tn5 enzyme (Illumina). The reaction was incubated at 37°C for 10 min with a gentle vortex every 5 min. The tagmentation reaction was inactivated by adding RIPA-low salt buffer. Crosslinks were reversed with Proteinase K digestion for 1 hr at 55°C then 65°C overnight. DNA was purified using a DNA Clean and Concentrator-5 kit (Zymo). ChIPmentation libraries were sequenced using 2 X 50 paired-end reads on an Illumina NextSeq 550. Sequencing data was processed as previously described (22).

**Lmx1b protein structure model**

The 3D structure of the *Drosophila* *engrailed* homeodomain (1HDD) was obtained from the RCSB PDB (23) <https://doi.org/10.1093/nar/28.1.235>.) and visualized with PyMOL (Schrödinger, LLC).

**Quantification and Statistical Analysis**
One-way ANOVA was performed for all analyses involving 3 or more genotypes using Welch’s correction where noted in the figure legend. An unpaired t-test with Welch’s correction was used for behavioral analyses. Graphpad Prism 8 was used for analysis and graphs using mean ± SEM. Statistical tests for gene list overlap or peak overlap were performed using a hypergeometric test (ChipPeakAnno, (24)) or a permutation test using 1000 permutations against randomly selected background regions (regioneR, (25). Pearson correlations were performed in R.

**Data and code availability**

All RNA-seq, ATAC-seq, and ChIPmentation datasets are available at the NCBI GEO (Gene Expression Omnibus) with accession identifiers GSE283961, GSE283962, GSE283963.

**Supplemental Figures**

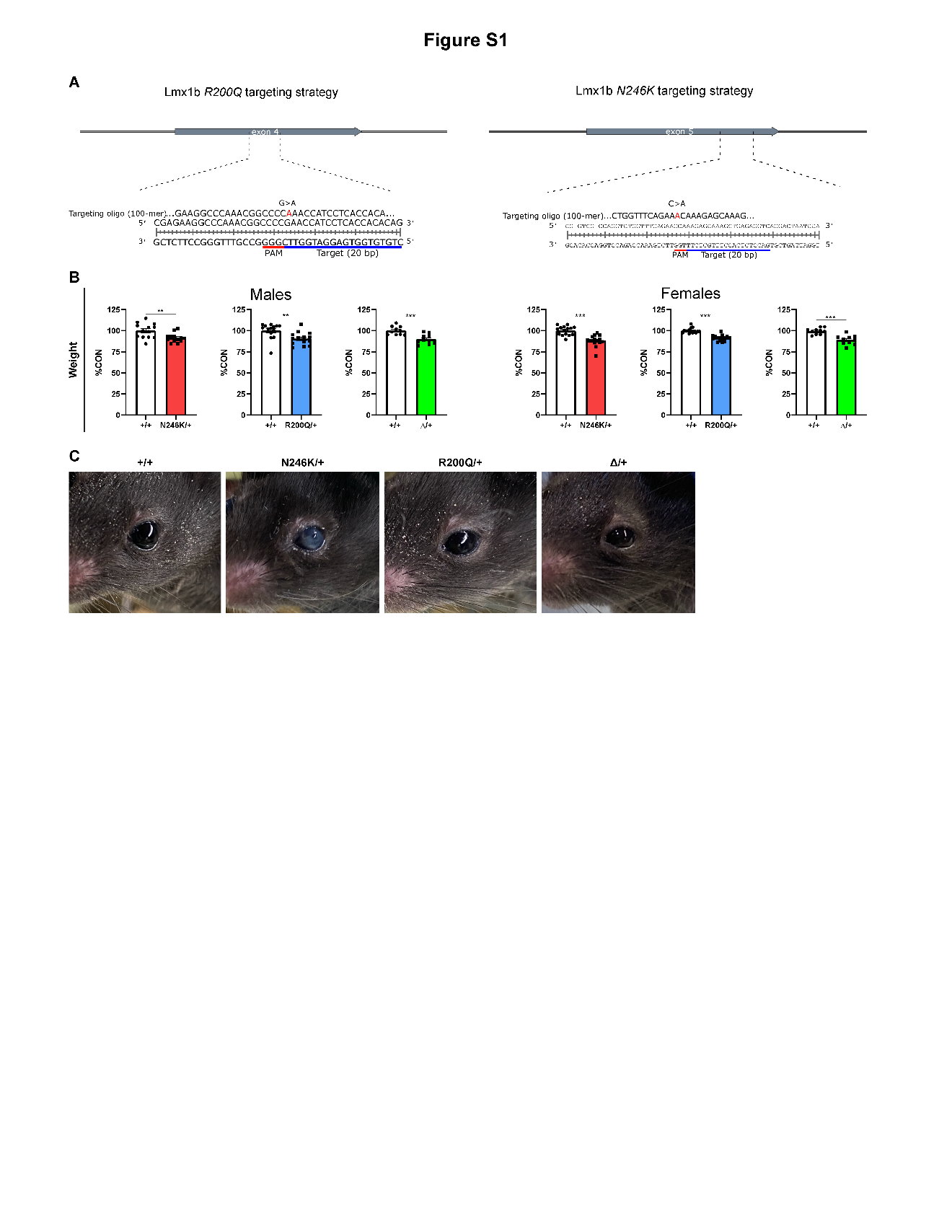

**Fig. S1: *Lmx1b* missense mouse lines**

A) Targeting strategy for introduction of *N246K* and *R200Q* point mutations.

B) *Lmx1b^N246K/+^*, *Lmx1b^R200Q^*^/+^and *Lmx1b^Δ/+^* adult body weights.

C) *Lmx1b^N246K/+^* eye pathology.

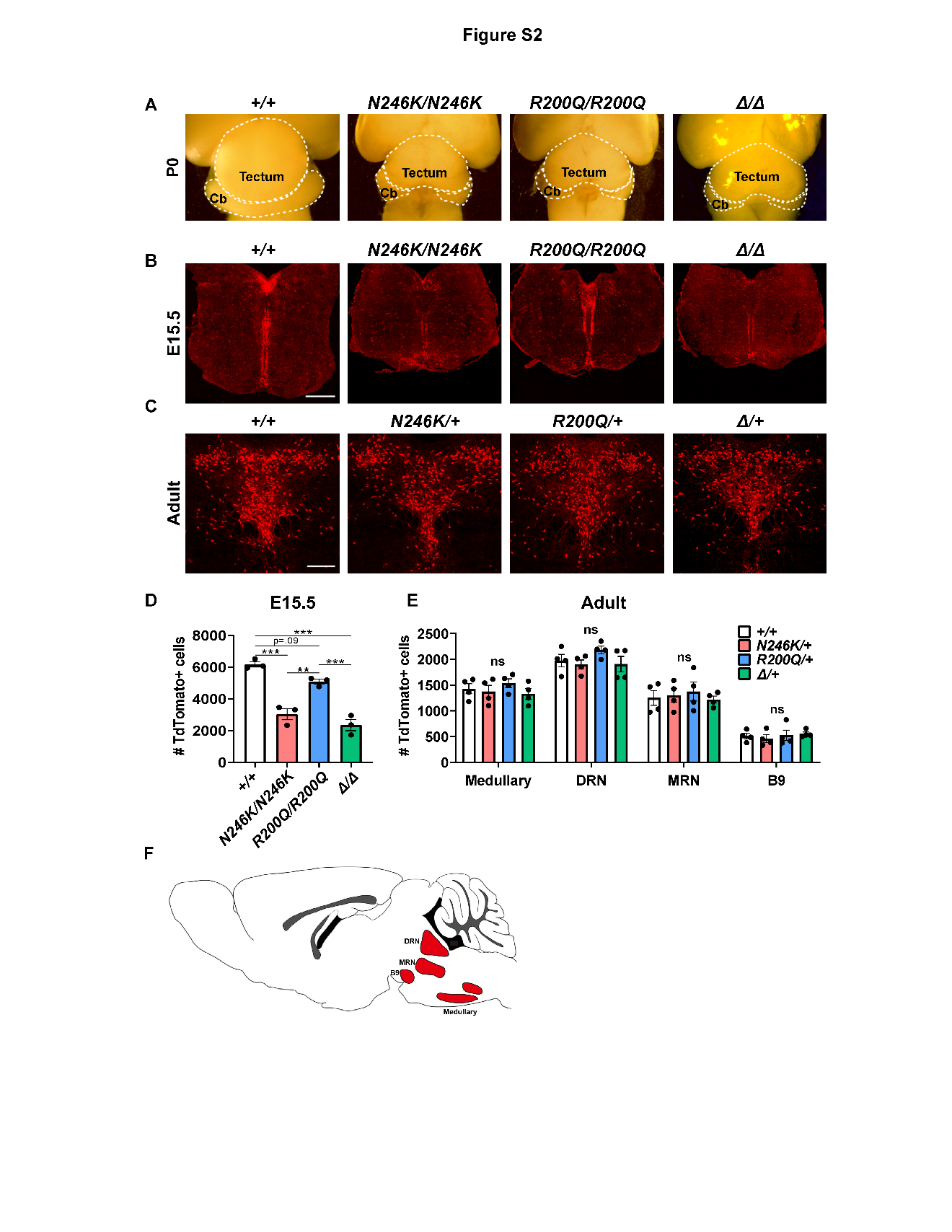

**Fig. S2: Homozygous and heterozygous missense brain morphology and *Pet1* neuron numbers**

A) Homozygous mutant brain morphology at birth (dorsal view). Cb, cerebellum. n=3 mice per genotype.

B) TdTomato^+^ (anti-RFP) *Pet1* neurons in E15.5 homozygous mutant brains. Scale bar, 500µm.

C) TdTomato^+^ (anti-RFP) *Pet1* (5-HT) neurons in adult heterozygous mutant dorsal raphe. Scale bar, 200µm.

D-E) TdTomato^+^ cells were counted across serially collected matched sections. ±SEM; n=3-4 mice per genotype; one-way ANOVA with Welch’s correction. **p<0.01, ***p<0.001, ns=not significant.

F) Adult raphe nuclei anatomical organization. *Pet1* neurons develop into mature 5-HT neurons (C) whose soma in the mammalian brain are located in the raphe nuclei. Axons issuing from 5-HT neurons in the DRN, MRN, and B9 project to the forebrain while axons from 5-HT cell bodies positioned in the medullary raphe nuclei project to the spinal cord. DRN, dorsal raphe nucleus; MRN, median raphe nucleus; B9, supralemniscal nucleus; Medullary, raphe nuclei in ventral medulla.

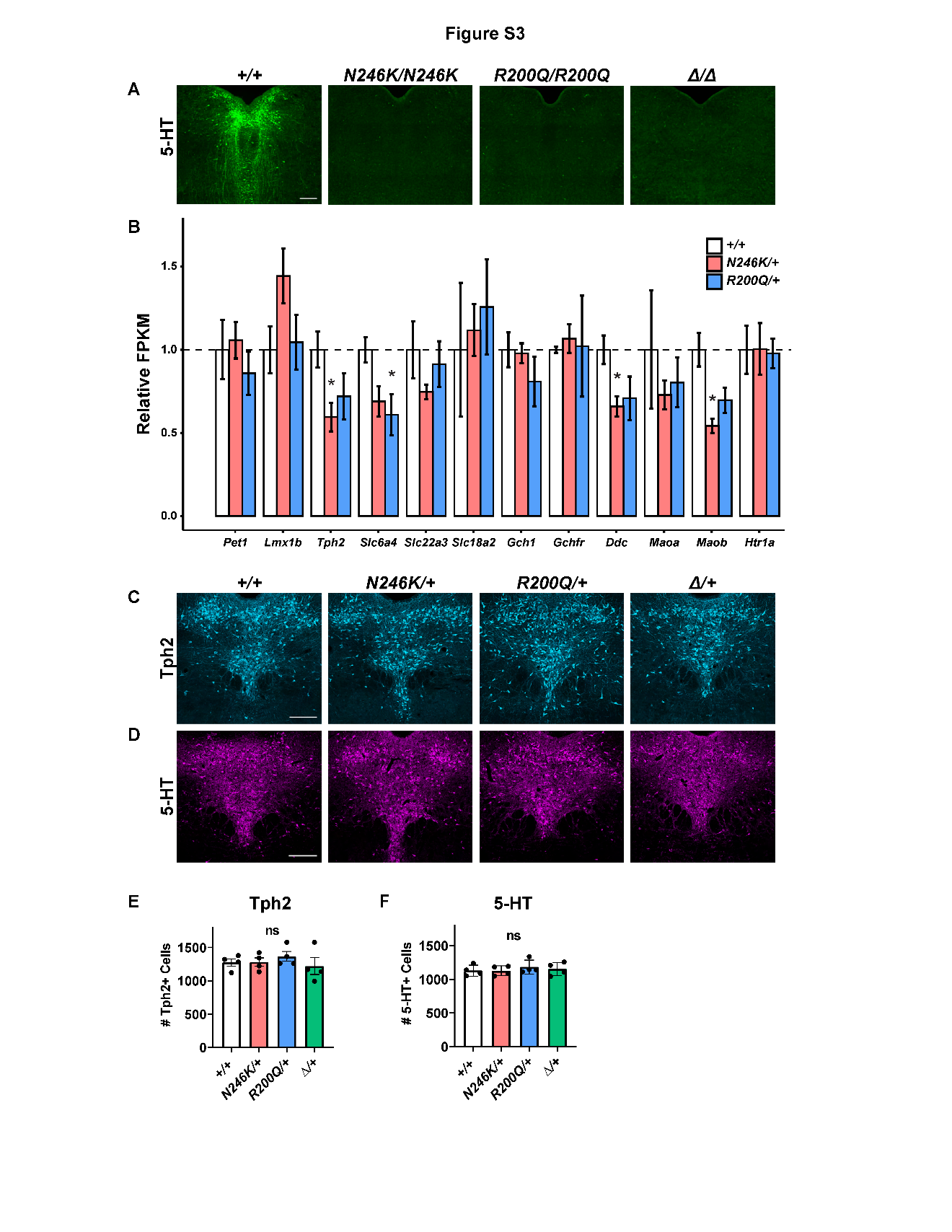

**Fig. S3: 5-HT identity in homozygous and heterozygous mutant *Pet1* neurons**

A) 5-HT immunostaining at E15.5 in homozygous missense and HD-deleted (Δ) mice. Scale bar, 200µm.

B) Relative expression levels (FPKM) of 5-HT neurotransmission genes at E17.5. *FDR≤0.01.

C) Representative coronal images of Tph2^+^ neurons in adult heterozygous mutant DRN. Scale bar, 200µm.

D) Representative coronal plane images of 5-HT^+^ neurons in adult heterozygous mutant DRN. Scale bar, 200µm.

E-F) Tph2^+^ (E) and 5-HT^+^ (F) neuron numbers in the DRN were estimated with serially collected matched sections. ±SEM; n=3-4 mice per genotype; one-way ANOVA with Welch’s correction. ns=not significant.

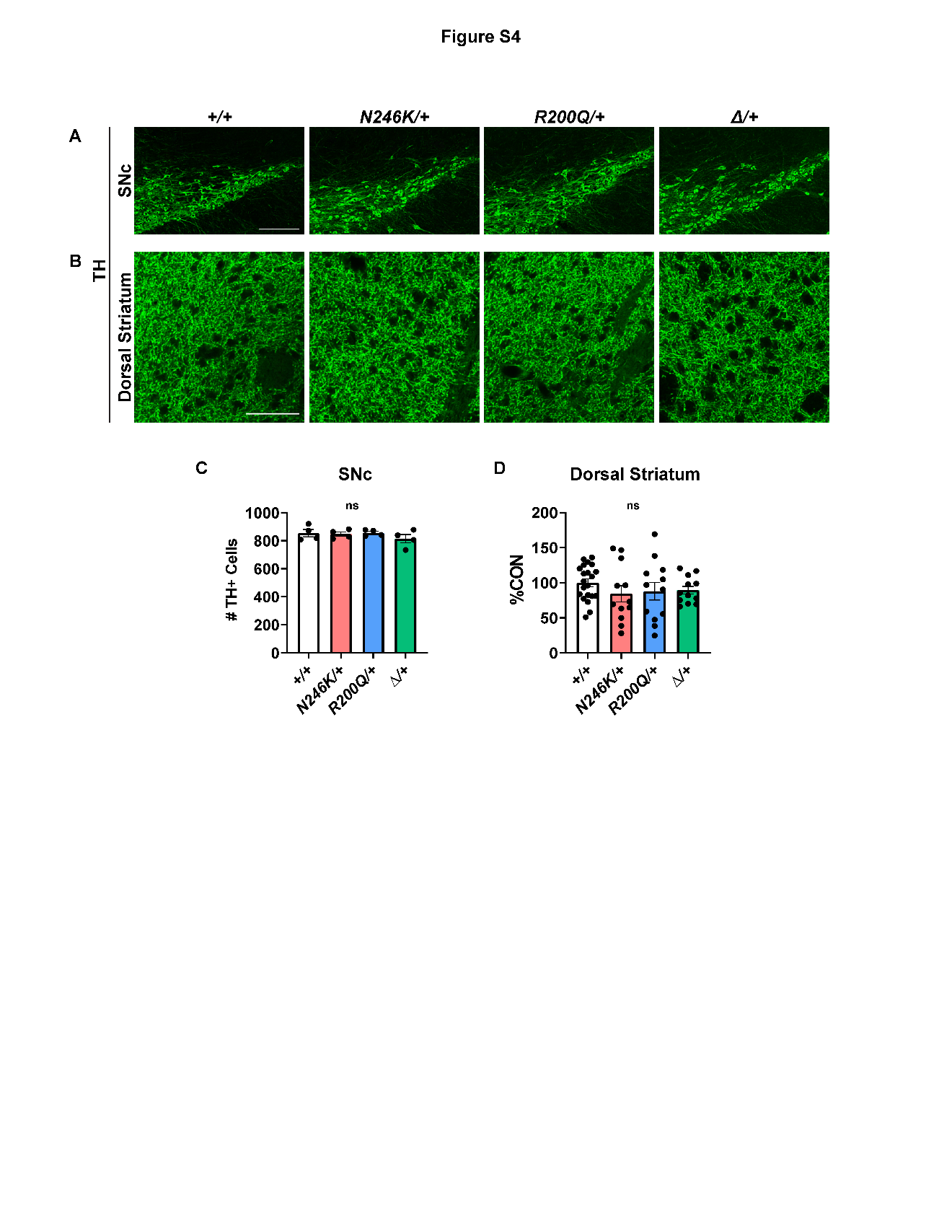

**Fig. S4: Dopamine neuron identity in adult heterozygous mutants**

A) Tyrosine hydroxylase (TH) immunostaining in substantia nigra pars compacta (SNc). Scale bar, 200µm.

B) TH immunostaining in the dorsal striatum. Scale bar, 50µm.

C) TH^+^ cell numbers in heterozygous mutant SNc from serially collected matched sections. n=4 mice per genotype; one-way ANOVA with Welch’s correction.

D) Relative TH^+^ pixel densities from three representative sections per biological replicate; ±SEM; n=4-7 mice per genotype; one-way ANOVA with Welch’s correction. ns=not significant.

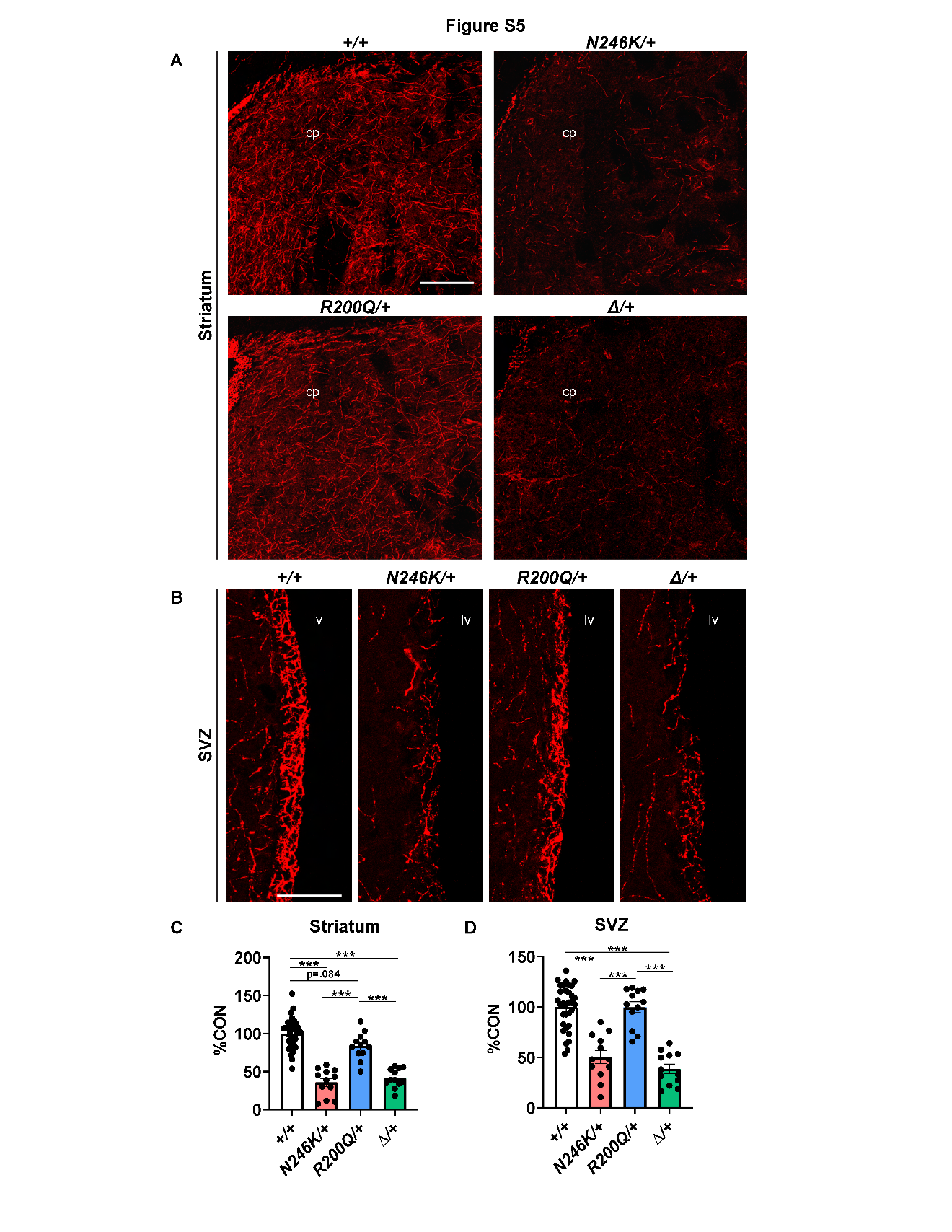

**Fig. S5: 5-HT axons in striatum and SVZ of adult heterozygous mutant mice.**

A) Representative images of TdTomato^+^ (anti-RFP) axons in the striatum. Scale bar, 100µm. cp, caudoputamen.

B) Representative images of TdTomato^+^ (anti-RFP) axons in the subventricular zone (SVZ). Scale bar, 50µm. lv, lateral ventricle.

C-D) Relative TdTomato^+^ (anti-RFP) pixel densities from three representative sections per biological replicate; ±SEM; n=4-11 mice per genotype; one-way ANOVA with Welch’s correction. ***p<0.001.

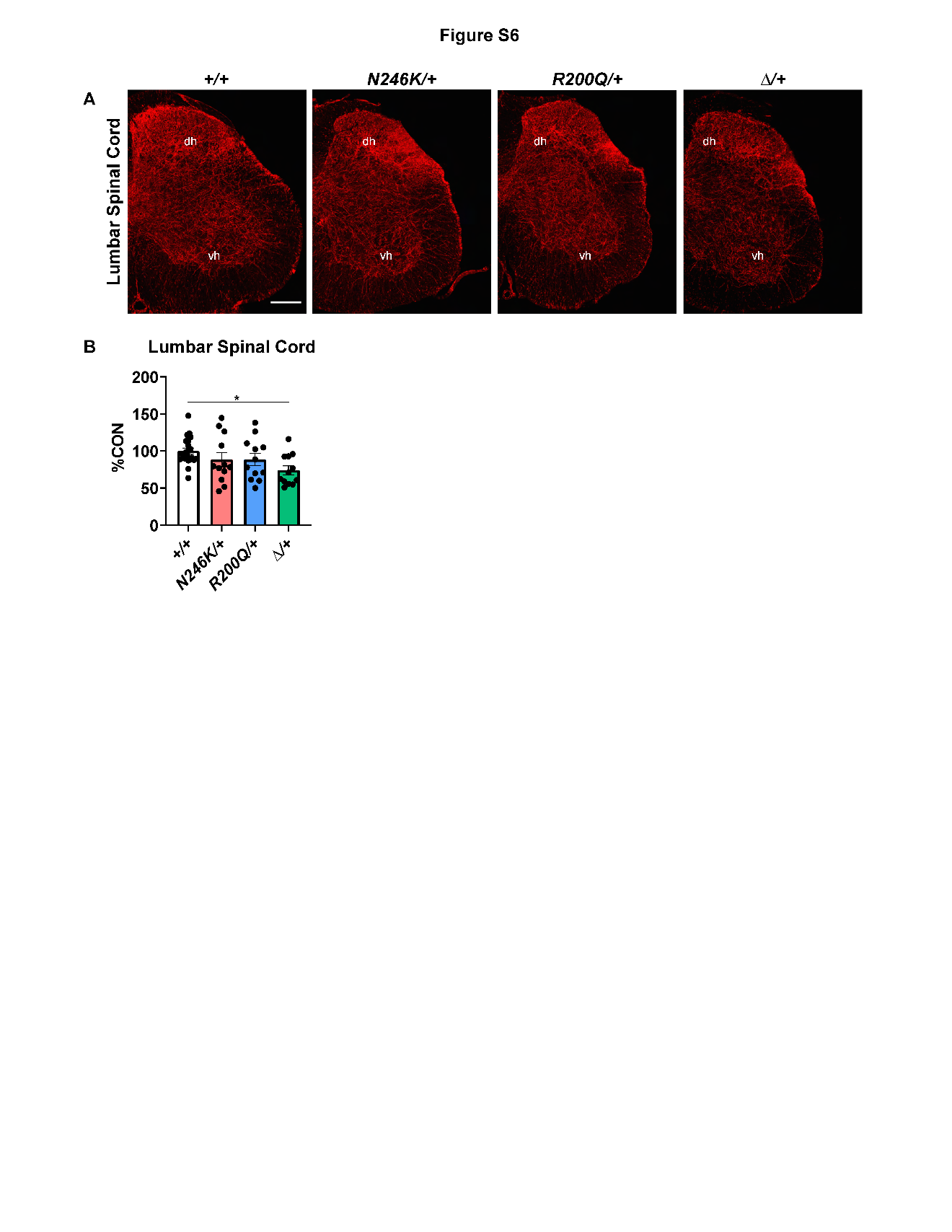

**Fig. S6: Axon densities in adult heterozygous mutant spinal cords**

A) Representative images of TdTomato^+^ axons (anti-RFP) in the lumbar spinal cord. Scale bar, 200µm. dh, dorsal horn. vh, ventral horn.

B) Relative TdTomato^+^ (anti-RFP) pixel densities from three representative sections per biological replicate; ±SEM; n=4-7 mice per genotype; one-way ANOVA with Welch’s correction. *p<0.05.

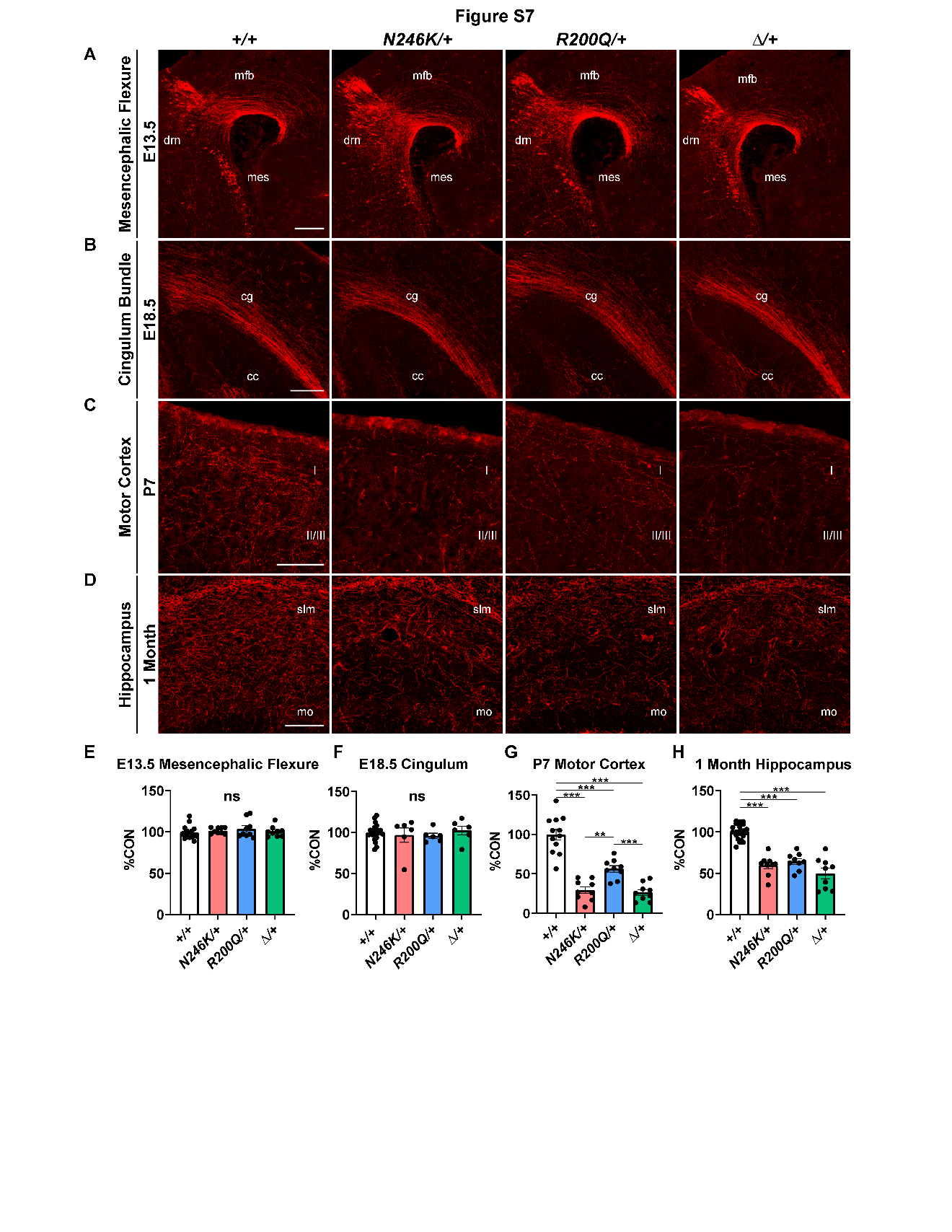

**Fig. S7: Postnatal 5-HT axon arbor formation is selectively disrupted in heterozygous missense mice**

A) Representative images of TdTomato^+^ (anti-RFP) axons in the mesencephalic flexure at E13.5. Scale bar, 200µm. drn, dorsal raphe nuclei; mfb, medial forebrain bundle; mes, mesencephalic flexure.

B) Representative images of TdTomato^+^ (anti-RFP) axons in the cingulum bundle at E18.5. Scale bar, 200µm. cg, cingulum bundle; cc, corpus callosum.

C) Representative images of TdTomato^+^ (anti-RFP) axons in the P7 motor cortex. Scale bar, 100µm.

D) Representative images of TdTomato^+^ (anti-RFP) axons in the hippocampus of 1 month-old mice. Scale bar, 100µm. slm, stratum lacunosum moleculare; mo, molecular layer.

E-H) Relative TdTomato^+^ (anti-RFP) pixel densities from 2-3 representative sections per biological replicate; ±SEM; n=3-9 mice per genotype; one-way ANOVA with Welch’s correction. ***p<0.001, ns=not significant.

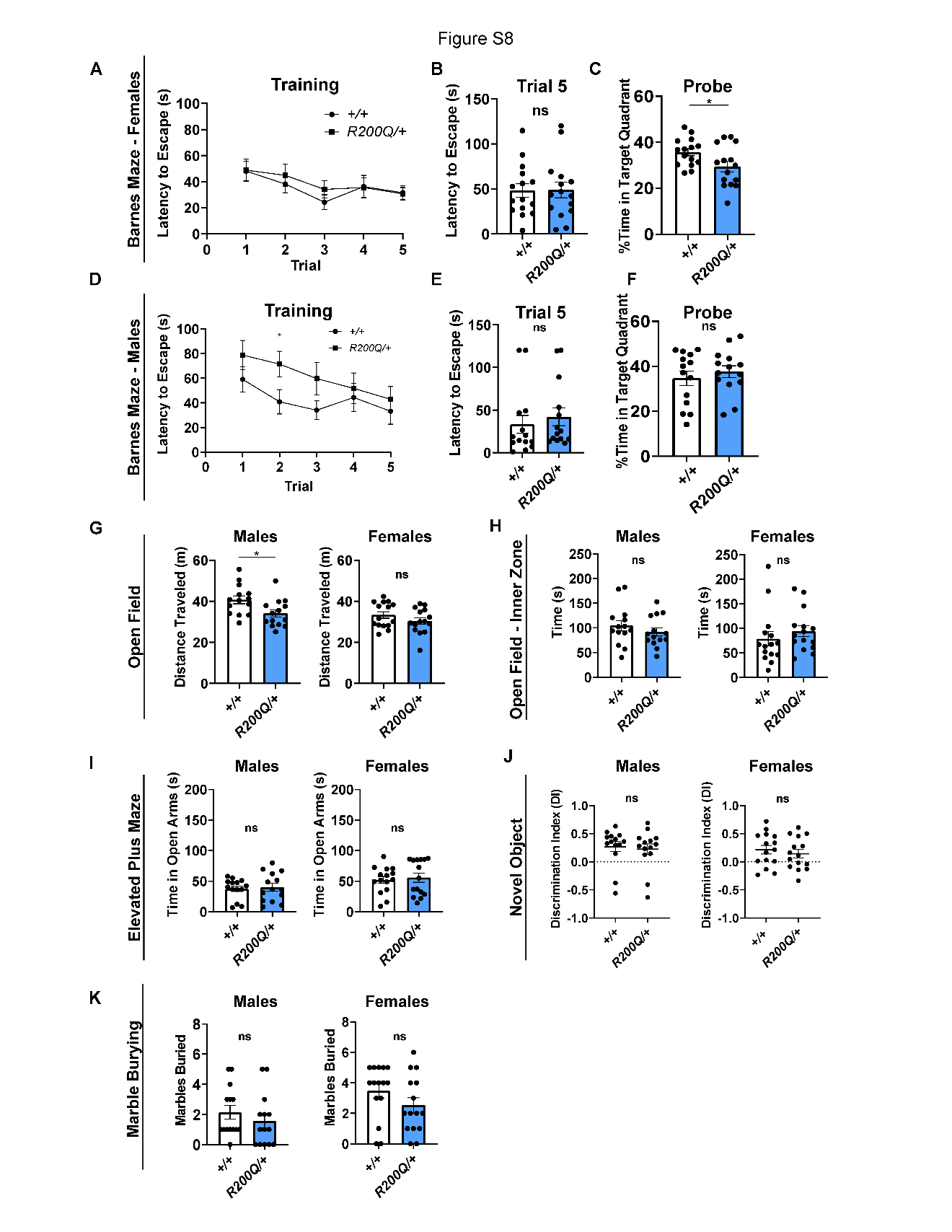

**Fig. S8: Behavior assays of *Lmx1b^R200Q/+^* mice.**

A-C) Barnes maze assay of 6–12-week-old females. A, Average latency to escape across five trials. B, Average escape latency in trial 5. C, Barnes maze probe trial. ±SEM; n=14-15 mice per genotype; Unpaired t-test with Welch’s correction.

D-F) Barnes maze assay of 6–12-week-old males. D, Average latency to escape across five trials. E, Average escape latency in trial 5. F, Barnes maze probe trial. ±SEM; n=15 mice per genotype; Unpaired t-test with Welch’s correction.

G) Open field, total distance traveled in 6–12-week-old mice. ±SEM; n=14-15 mice per genotype; Unpaired t-test with Welch’s correction.

H) Open field, total time spent in the inner zone of the arena in 6–12-week-old mice. ±SEM; n=14-15 mice per genotype; Unpaired t-test with Welch’s correction.

I) Elevated plus maze. Total time spent in the open arms of the maze in 6–12-week-old mice. ±SEM; n=14-15 mice per genotype; Unpaired t-test with Welch’s correction.

J) Novel object recognition in 6–12-week-old mice. Discrimination index (DI): (Time spent exploring novel object – Time spent exploring familiar object)/(Total exploration time). ±SEM; n=14-15 mice per genotype; Unpaired t-test with Welch’s correction.

K) Marble burying. Total marbles buried in 6–12-week-old mice. ±SEM; n=14-15 mice per genotype; Unpaired t-test with Welch’s correction.

*p<0.05, ns=not significant.

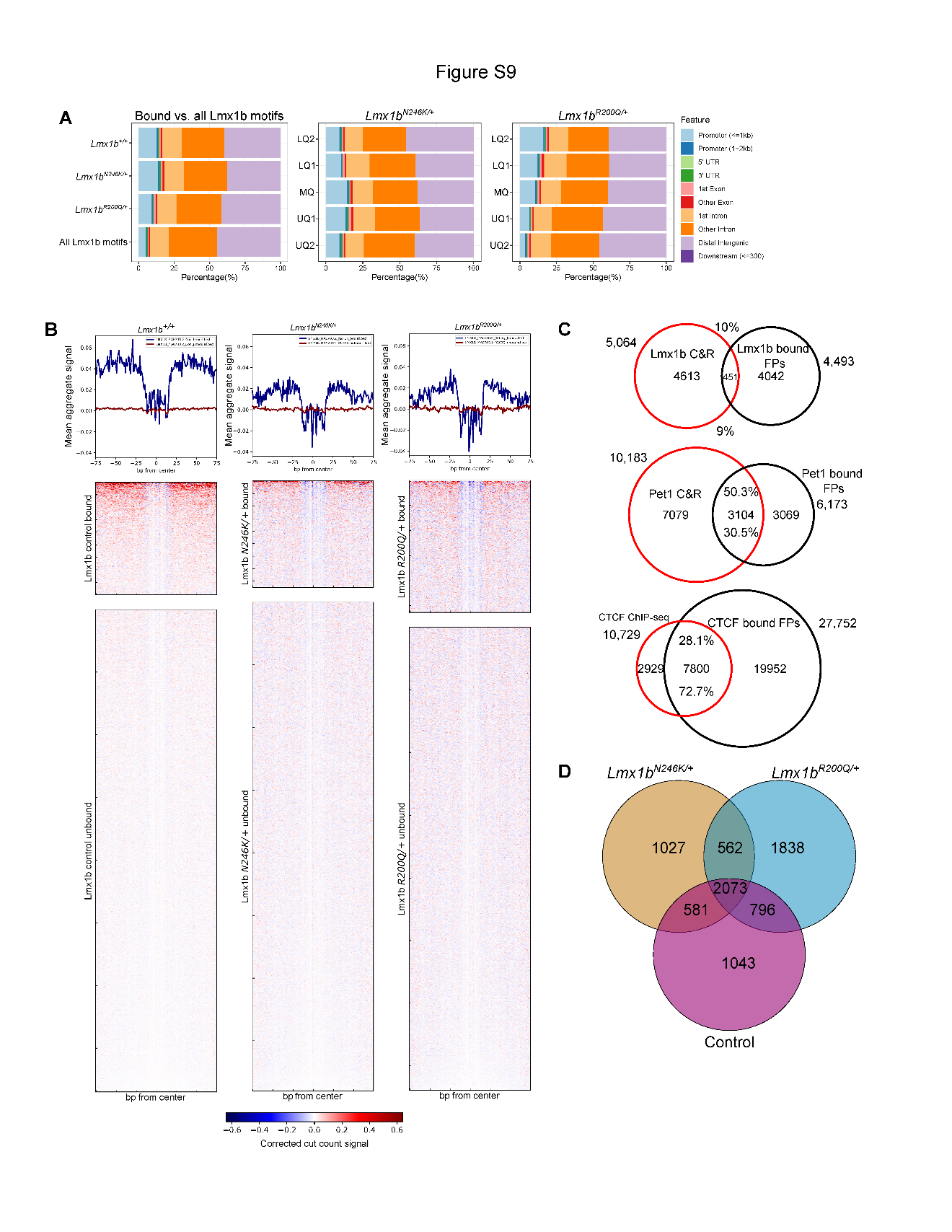

**Fig. S9: Lmx1b footprinting and intersection with TF CUT&RUN peaks**

A) Distribution of bound Lmx1b footprints in *Lmx1b^+/+^*, *Lmx1b^N246K/+^*, and *Lmx1b^R200Q^*^/+^ *Pet1* neurons vs all Lmx1b motifs.

B) Individual aggregate Lmx1b motif footprint profiles and bound and unbound heatmaps in *Lmx1b^+/+^*, *Lmx1b^N246K/+^*, and *Lmx1b^R200Q^*^/+^ *Pet1* neurons. Heatmap color scale: red, higher than expected cut counts; blue, lower than expected cut counts.

C) Bound Lmx1b motif overlaps in *Lmx1b^+/+^*, *Lmx1b^N246K/+^*, and *Lmx1b^R200Q^*.

D) Overlaps between bound footprint regions for Lmx1b, Pet1, and CTCF with Lmx1b CUT&RUN, Pet1 CUT&RUN, and CTCF ChIPmentation, respectively. Permutation test p-value < 0.001.

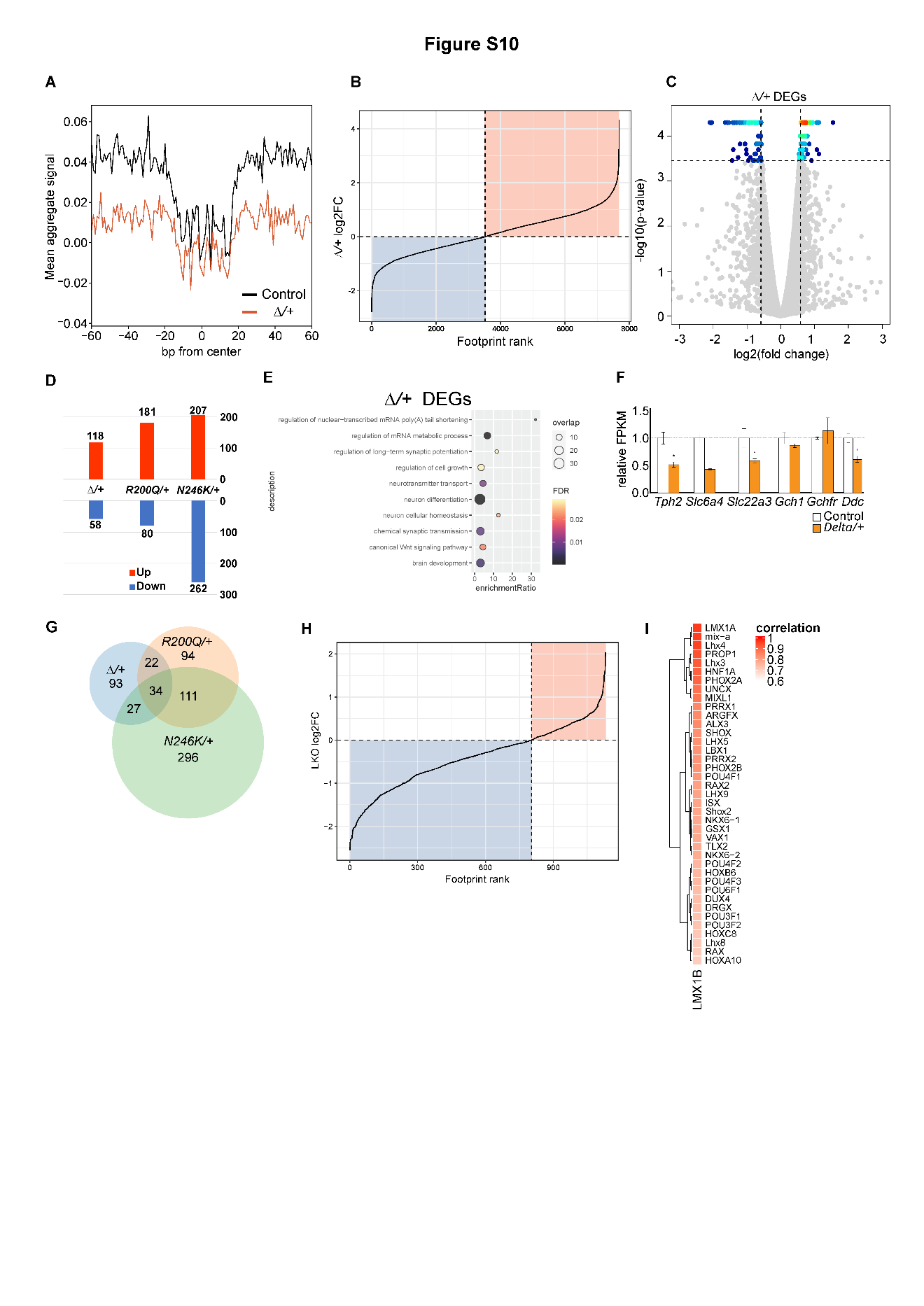

**Fig. S10: *Lmx1b Δ* allele DGF and gene expression**

A) Aggregate footprint profile for Lmx1b motif binding activity in *Lmx1b^Δ/+^* (orange) *Pet1* neurons vs. control (black) *Pet1* neurons.

B) Log2FC plot of Lmx1b motif footprint scores in *Lmx1b^Δ/+^* *Pet1* neurons. Colored quadrants highlight proportion of footprint losses (blue) and footprint gains (red). Note greater proportion of gains similar to *R200Q/+.*

C) *Lmx1b^Δ/+^* RNA-seq volcano plot. Colored dots indicate significant DEGs. Dot heatmap indicates dot density: red, high; yellow/cyan, intermediate; blue, low. FC ≥ 1.5 and FDR < 0.01 shown as -log10 p-value.

D) *Lmx1b^Δ/+^*, *Lmx1b^N246K/+^*, and *Lmx1b^R200Q^* up and down DEG numbers.

E) *Lmx1b^Δ/+^* DEG GO term enrichment.

F) Relative expression levels (FPKM) of 5-HT neurotransmission genes at E17.5 in *Lmx1b^Δ/+^ Pet1* neurons. *FDR≤0.01.

G) *Lmx1b^Δ/+^*, *Lmx1b^N246K/+^* and *Lmx1b^R200Q^*^/+^ DEG overlaps.

H) Log2FC plot of Lmx1b motif footprint scores in *Lmx1b^f/f;Pet1-Cre;Ai9^* (LKO) *Pet1* neurons. ATAC-seq libraries were not sequenced as deeply as those for mutant heterozygous libraries thus resulting in fewer overall numbers of footprints (x-axis).

I) HD motif similarity heatmap (Pearson correlation coefficient, p<0.01).

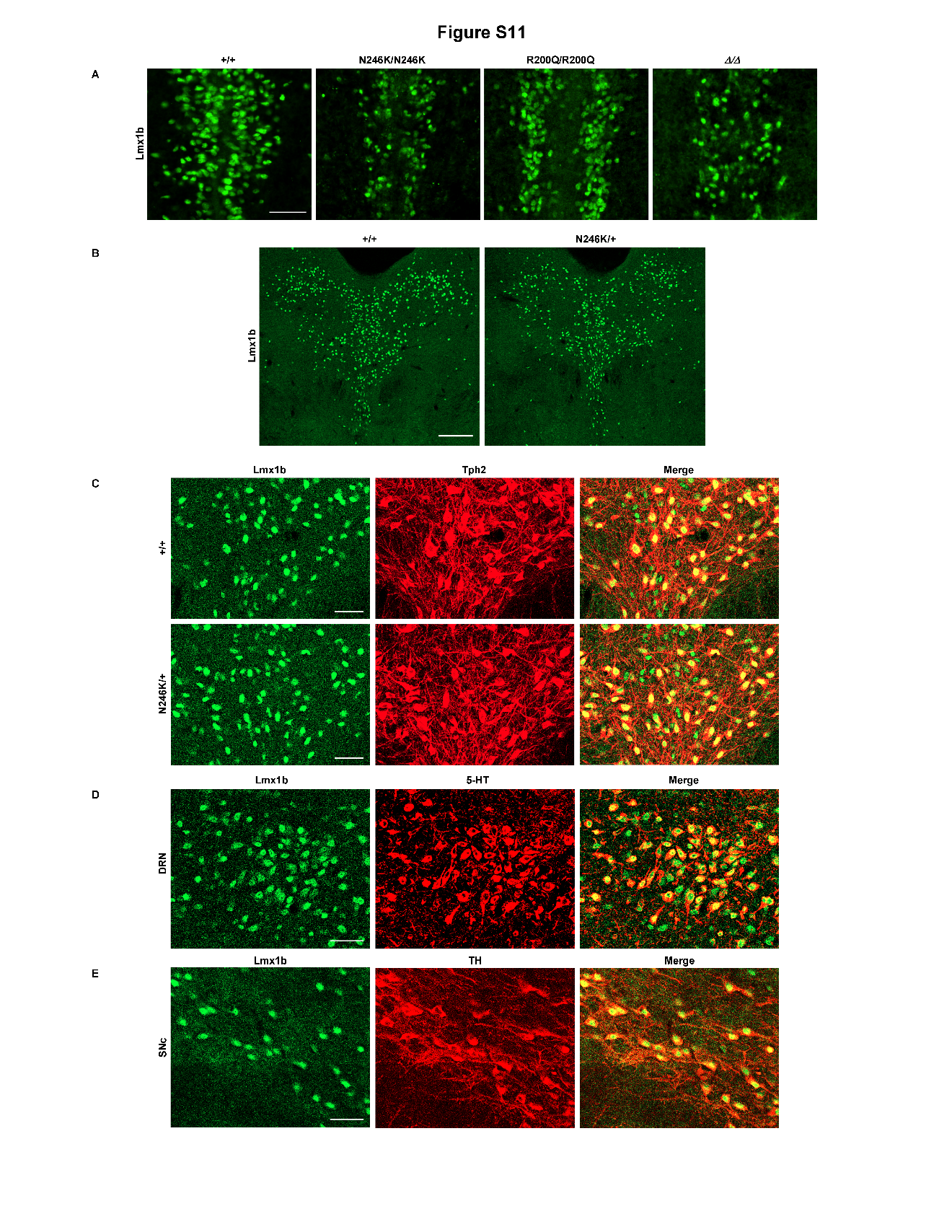

**Fig. S11: Missense protein expression**

A) Lmx1b immunostaining in rostral hindbrain at E15.5 in *Lmx1b^+/+^*, *Lmx1b^N246K/N246K^*, *Lmx1b^R200Q/R200Q^* and *Lmx1b^Δ/Δ^* mice. Scale bar, 50 µm.

B) Lmx1b immunostaining in control (+/+) and *Lmx1b^N246K/+^* adult DRN.

C) Lmx1b and Tph2 co-immunostaining in adult DRN. Scale bar, 50 µm.

D) Lmx1b and 5-HT co-immunostaining in control adult DRN. Scale bar, 50 µm.

E) Lmx1b and TH co-immunostaining in control adult SNc.

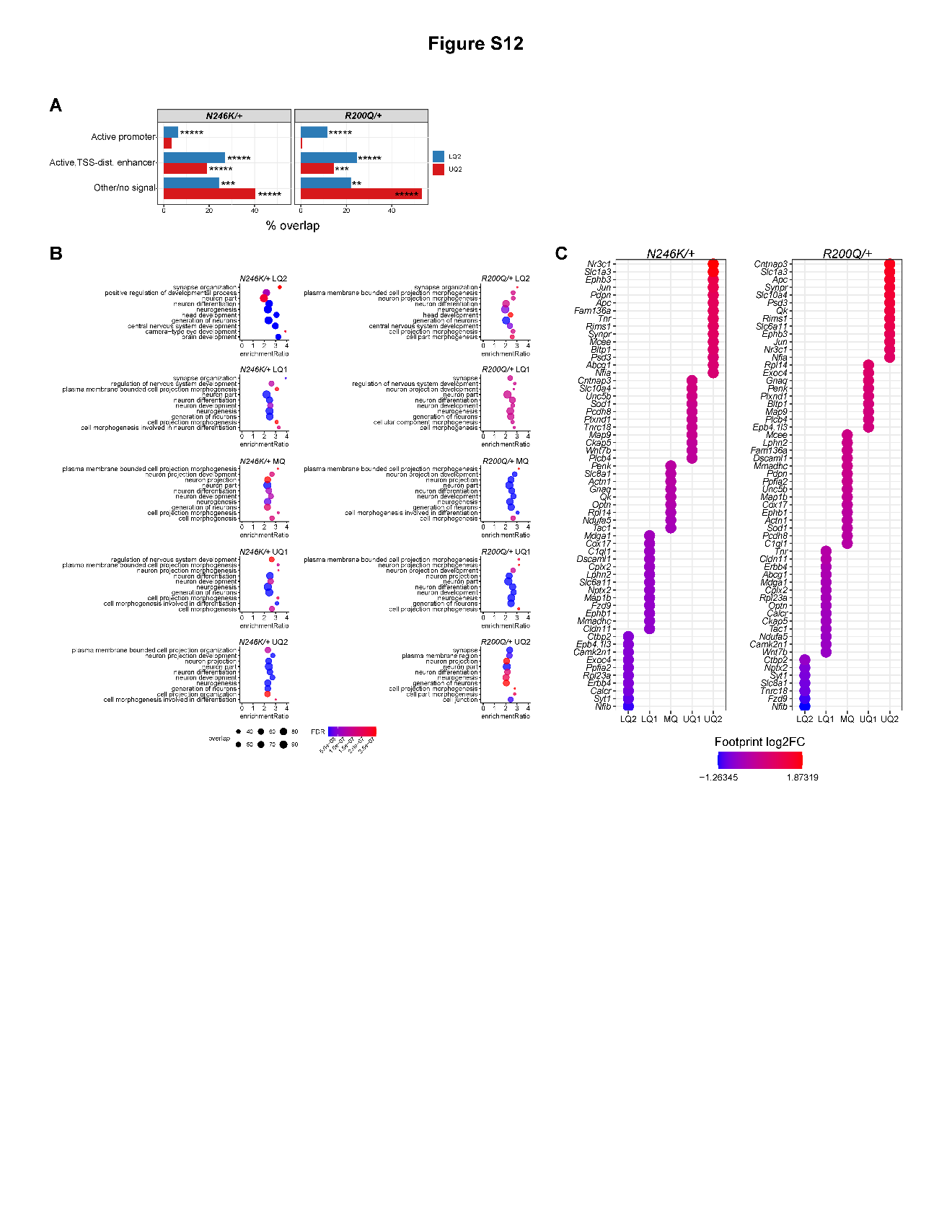

**Fig. S12: Lmx1b footprint chromatin states, GO term enrichment and association to SAM DEGs**

A) Chromatin state comparison of LQ2 vs UQ2 motifs in *Lmx1b^N246K/+^* and *Lmx1b^R200Q^*^/+^. Hypergeometric test p-values, ***** ≤ 0.00001, *** ≤ 0.001, ** ≤ 0.01.

B) Gene ontology enrichment analysis of genes associated with Lmx1b footprint score changes in each quintile in *Lmx1b^N246K/+^* (left) and *Lmx1b^R200Q^*^/+^ (right) *Pet1* neurons.

C) Plot of footprint score changes across quintiles associated with common SAM DEGs in *Lmx1b^N246K/+^* (left) and *Lmx1b^R200Q/+^* (right) *Pet1* neurons. Dot colors represent log2FC of individual Lmx1b motif footprints and grouped accordingly into quintiles.

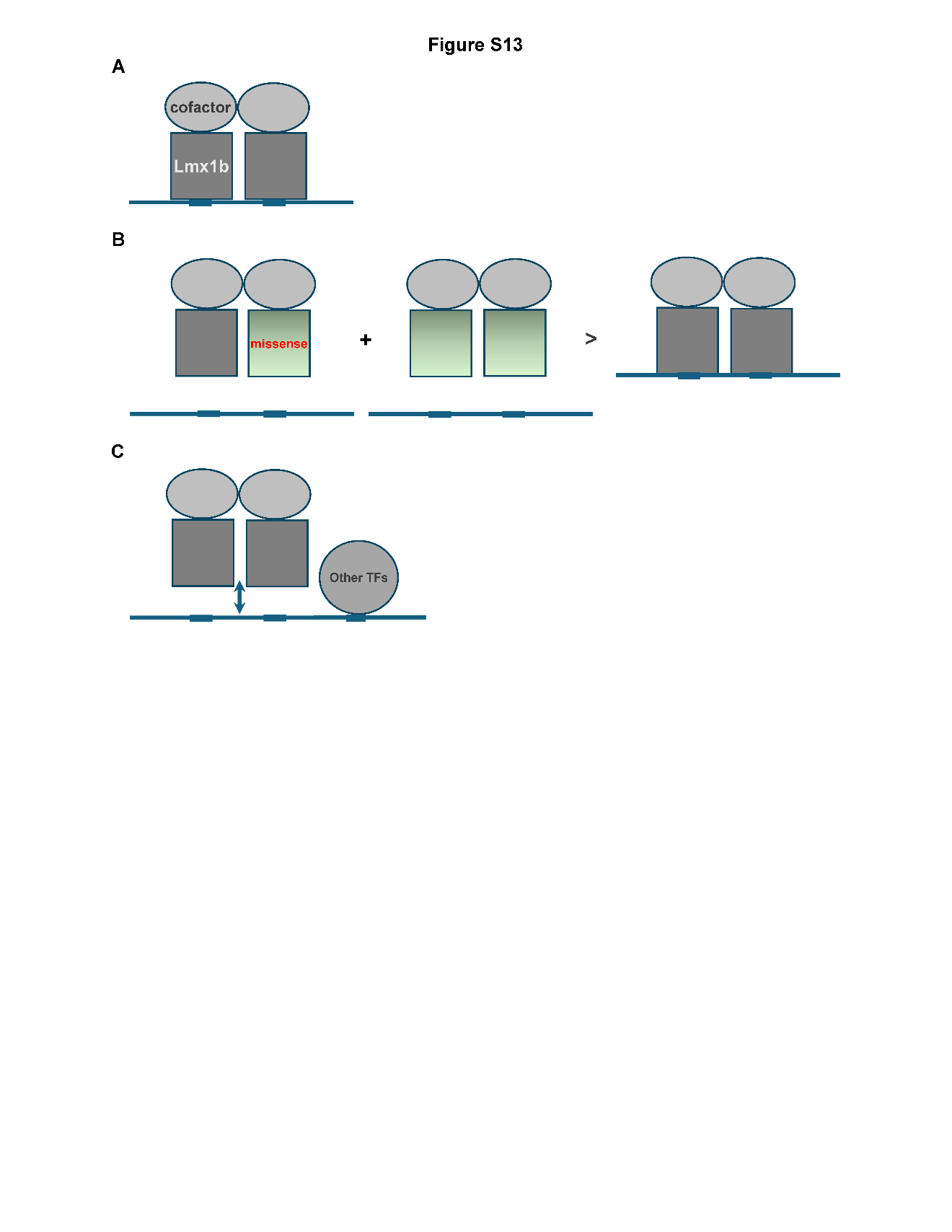

**Fig. S13: Interpretation of DGF findings**

A. Lmx1b binding to Lmx1b motifs (gray squares) depends on complex formation with cofactors (ovals) such as LIM domain binding proteins or perhaps other sequence-specific TFs (52, 53).

B. In mutant heterozygous *Pet1* neurons, mutant Lmx1b (green squares) cannot bind Lmx1b motifs but still forms complexes that are nonfunctional and predominant over wildtype complexes thus interfering (antimorphic antagonism) with wildtype. Interference is sufficient for complete footprint loss at some motifs (LQ2). At other motifs residual functional Lmx1b complexes may be sufficient for partial (LQ1) or full (MQ) footprints.

C. In mutant heterozygous *Pet1* neurons, other HD TFs (gray circle) may bind with or without (bi-arrow) residual functional Lmx1b complexes to form new footprints or enhance existing ones (UQ1, UQ2).

Dataset S1 (separate file). List of *Lmx1b^N246K/+^*, *Lmx1b^R200Q^*^/+^ and *Lmx1b^Δ/+^* differentially expressed genes.
